# CDK11 Promotes Centromeric Transcription to Maintain Centromeric Cohesion during Mitosis

**DOI:** 10.1101/2022.02.01.478617

**Authors:** Qian Zhang, Yujue Chen, Zhen Teng, Zhen Lin, Hong Liu

**Affiliations:** Department of Biochemistry and Molecular Biology, Tulane University School of Medicine, 1430 Tulane Ave, New Orleans, LA 70112, USA; Tulane Cancer Center, Tulane University School of Medicine, New Orleans, LA, 70112, USA; Tulane Aging Center, Tulane University School of Medicine, New Orleans, LA, 70112, USA; Department of Pathology and Laboratory Medicine, Tulane University School of Medicine, 1430 Tulane Ave, New Orleans, LA 70112, USA

## Abstract

Actively-transcribing RNA polymerase (RNAP)II is remained on centromeres to maintain centromeric cohesion during mitosis although it is largely released from chromosome arms. This pool of RNAPII plays an important role in centromere functions. However, the mechanism of RNAPII retention on mitotic centromeres is poorly understood. We here demonstrate that Cdk11 depletion-induced centromeric cohesion defects are largely independent of Bub1. We further show that Cdk11 depletion and expression of its kinase-dead version significantly reduce both RNAPII and elongating RNAPII (pSer2) levels on centromeres, and also decrease centromeric transcription without altering the protein expression of cohesin and cohesion-regulators. Interestingly, enhanced centromeric transcription by THZ1 treatment or overexpression of CENP-B DNA-binding domain completely rescues Cdk11-depletion defects. These results suggest that Cdk11 promotes centromeric cohesion through facilitating centromeric transcription. Mechanistically, Cdk11 binds and phosphorylates RNAPII to promote transcription. Furthermore, mitosis-specific degradation of G2/M Cdk11-p58 recapitulates Cdk11-depletion defects. Altogether, our findings establish Cdk11 as an important regulator of centromeric transcription and as part of the mechanism for retaining RNAPII on centromeres during mitosis.

## Introduction

The non-coding centromere, a specialized region of a chromosome, dictates the assembly of kinetochore that is essential for proper chromosome segregation during mitosis. Such a critical centromere function is conserved across eukaryotes and is determined by the centromere-specific histone H3 variant CENP-A (McKinley and Cheeseman, 2016). Proper incorporation of CENP-A into centromeric chromatin is prerequisite for CENP-A to fulfill its duty. Unlike canonical histones that are usually incorporated into chromatin in a DNA duplication-dependent manner during S phase, newly synthesized CENP-A is instead deposited into centromeric chromatin independently of DNA replication mainly during G1 phase (Jansen et al., 2007; Schuh et al., 2007). This process requires RNA polymerase (RNAP)II-catalyzed transcription (Bobkov et al., 2018; Bury et al., 2020), suggestive of an essential role of centromeric transcription in centromere functions. In addition to in interphase, centromeric transcription is also undergoing during mitosis (Bobkov et al., 2018; Chan et al., 2012; Chen et al., 2021; Liu et al., 2015; Perea-Resa et al., 2020). The ongoing centromeric transcription facilitates the installment of an essential cohesion-protector Sgo1 onto centromeres to protect centromeric cohesion during mitosis in human cells (Chen et al., 2021; Liu et al., 2015). Thus, centromeric transcription plays diverse critical roles in regulating centromere functions and chromosome segregation (Talbert and Henikoff, 2018).

As such, how centromeric transcription is regulated is poorly understood. In budding yeast, centromere-binding factor cbf1 and histone H2A variant Htz1 have been demonstrated to repress centromeric transcription (Ling and Yuen, 2019). In human cells, the nucleolus and ZFAT was suggested to regulate centromeric transcription as well (Bury et al., 2020; Ishikura et al., 2020). Thus, there exist factors that may be more specific for centromeric transcription. Identification of these factors would be a great help of understanding the regulation of centromeric transcription. When cells enter mitosis, RNAPII and its associated factors are largely released from chromosomes, thus leading to a global suppression for transcription (Palozola et al., 2018; Parsons and Spencer, 1997; Teves et al., 2018). However, a pool of RNAPII is still remained on mitotic centromeres to actively transcribe centromeres (Bobkov et al., 2018; Chan et al., 2012; Chen et al., 2021; Liu et al., 2015; Perea-Resa et al., 2020). Therefore, a mechanism must exist to maintain active RNAPII on centromeres during mitosis. Identification of the factors specific to centromeric transcription during mitosis would help resolve such a mechanism.

Cdk11 belongs to the cyclin-dependent kinase family, which contains several members that are important for transcriptional regulation, including Cdk7 and Cdk9 (Chou et al., 2020). These kinases phosphorylate either RNAPII at Ser5 to facilitate transcriptional initiation or RNAPII at Ser2 to promote transcriptional elongation, establishing these kinases as universal regulators for RNAPII gene transcription (Fu et al., 1999; Lu et al., 1992; Peng et al., 1998; Serizawa et al., 1995; Spangler et al., 2001). Interestingly, extensive studies have suggested important roles of Cdk11 in transcriptional regulation. Firstly, Cdk11 regulates mRNA splicing and processing (Dickinson et al., 2002; Hu et al., 2003; Loyer et al., 2008; Loyer et al., 1998; Pak et al., 2015; Trembley et al., 2002; Valente et al., 2009). Secondly, in fission yeast, Cdk11 has been shown to phosphorylate the transcriptional mediator complex (Drogat et al., 2012). In addition, Cdk11 can also directly phosphorylate the Ser2 of the RNAPII CTD to regulate HIV viral transcription (Pak et al., 2015). More recently, Gajduskova *et al*. showed that Cdk11 promotes transcription of replication-dependent histone genes via phosphorylating Ser2 of the RNAPII CTD (Gajduskova et al., 2020). These observations suggest that, unlike the universal transcriptional regulators Cdk7 and Cdk9, Cdk11 may function exclusively on a subset of genes to enhance their transcription, which might be important for some specific cellular needs. As such, it would be of interest to determine whether Cdk11 could be important for the transcription of non-coding DNA sequences, such as centromeres. Several previous observations suggested that Cdk11 is involved in centromere regulation. Cdk11 knockout in mice resulted in early embryonic lethality due to apoptosis of the blastocyst cells (Li et al., 2004). Cells within these embryos exhibited mitotic arrest (Li et al., 2004). Cdk11 depletion in human cells also caused increase in mitosis-arrested cells that suffered severe centromeric cohesion defects (Hu et al., 2007; Rakkaa et al., 2014b). All these findings highlight the importance of Cdk11 in regulating centromeric cohesion. Interestingly, a Cdk11 isoform p58 is generated from an internal ribosomal site on Cdk11 mRNAs exclusively during G2/M phase (Cornelis et al., 2000; Xiang et al., 1994), suggesting a potential important role of Cdk11-p58 in mitotic regulation. Based on these observations, together with our findings that centromeric transcription facilitates centromeric cohesion (Chen et al., 2021), we thereby hypothesized that Cdk11 might promote centromeric transcription to maintain centromeric cohesion.

In this study, we address whether and how Cdk11 regulates centromeric transcription in human cells. Cdk11 depletion significantly reduces RNAPII and RNAPII-pSer2 levels at centromeric chromatin, reduces ongoing centromeric transcription and weakens centromeric cohesion. Enhanced centromeric transcription completely rescues Cdk11-depletion phenotypes. Importantly, mitosis-specific degradation of G2/M Cdk11 p58 recapitulates Cdk11-depletion defects. Thus, our findings establish Cdk11 as an important regulator of centromeric transcription as well as part of the mechanism for retaining RNAPII on centromeres during mitosis.

## Results

### Cdk11 depletion slightly decreases Bub1 recruitment to kinetochores

Cdk11 depletion resulted in centromeric cohesion defects in mitosis and the defects had been attributed to decreased Bub1 recruitment to kinetochores (Hu et al., 2007; Rakkaa et al., 2014a). However, the cohesion defects caused by Bub1 depletion per se seem much milder than those by Cdk11 (Hu et al., 2007; Liu et al., 2013a; Rakkaa et al., 2014b; Zhang et al., 2014), challenging the notion that decreased Bub1 levels on kinetochores is the major factor contributing to Cdk11 depletion-induced cohesion defects. Therefore, we re-examined Bub1 kinetochore localization in Cdk11 depletion cells. HeLa Tet-On cells transfected with luciferase (mock) or two distinct Cdk11 siRNA oligos were transiently treated with nocodazole and collected mitotic cells were subject to chromosome spread and immunostaining. As a comparison, Bub1 siRNA oligos were also included. Consistently, Bub1 levels on kinetochores were decreased by more than 90% upon Bub1 depletion, whereas they were only moderately reduced by ∼40% upon Cdk11 depletion (**Figures 1A** and **1B**). ∼ 90% of Cdk11 depletion cells exhibited impaired centromeric cohesion. Further analyses demonstrated a heterogeneity of Bub1 kinetochore localizations among these cells (**Figure 1A**). While Bub1 levels were reduced at kinetochores in 47% of cells after Cdk11 depletion, 32% of Cdk11-depleted cells still showed strong Bub1 signals with weakened centromeric cohesion and 11% of Cdk11-depleted cells showed strong Bub1 signals with severely impaired centromeric cohesion (**Figure 1A**). Thus, no obvious correlation between Bub1 kinetochore levels and the robustness of centromeric cohesion was observed. Importantly, with such a milder decrease in Bub1 levels on kinetochores, Cdk11 depletion even induced severer centromeric cohesion defects than Bub1 depletion, revealed by the sister-centromeres distance (**Figures 1B** and **1C**). These results strongly suggest that Bub1 is unlikely the major factor contributing to Cdk11-regulated centromeric cohesion. Bub1 recruitment to kinetochores is dependent on the phosphorylation of Knl1 MELT domains (Primorac et al., 2013; Zhang et al., 2014). To determine how Cdk11 depletion decreased Bub1 recruitment to kinetochores, we analyzed Knl1 phospho-MELT levels on kinetochores in Cdk11-depleted cells. HeLa Tet-On cells were transfected with luciferase (mock) or Cdk11 siRNAs. Cells were then treated with nocodazole for 2 hr and mitotic cells were collected for chromosome spread and immunostaining. We found that Cdk11 depletion indeed decreased the levels of Knl1 phospho-MELT by ∼40% (**Figures 1D** and **1E**) on kinetochores, likely explaining why Bub1 kinetochore recruitment is impaired in Cdk11 depletion cells.

**Figure 1.**
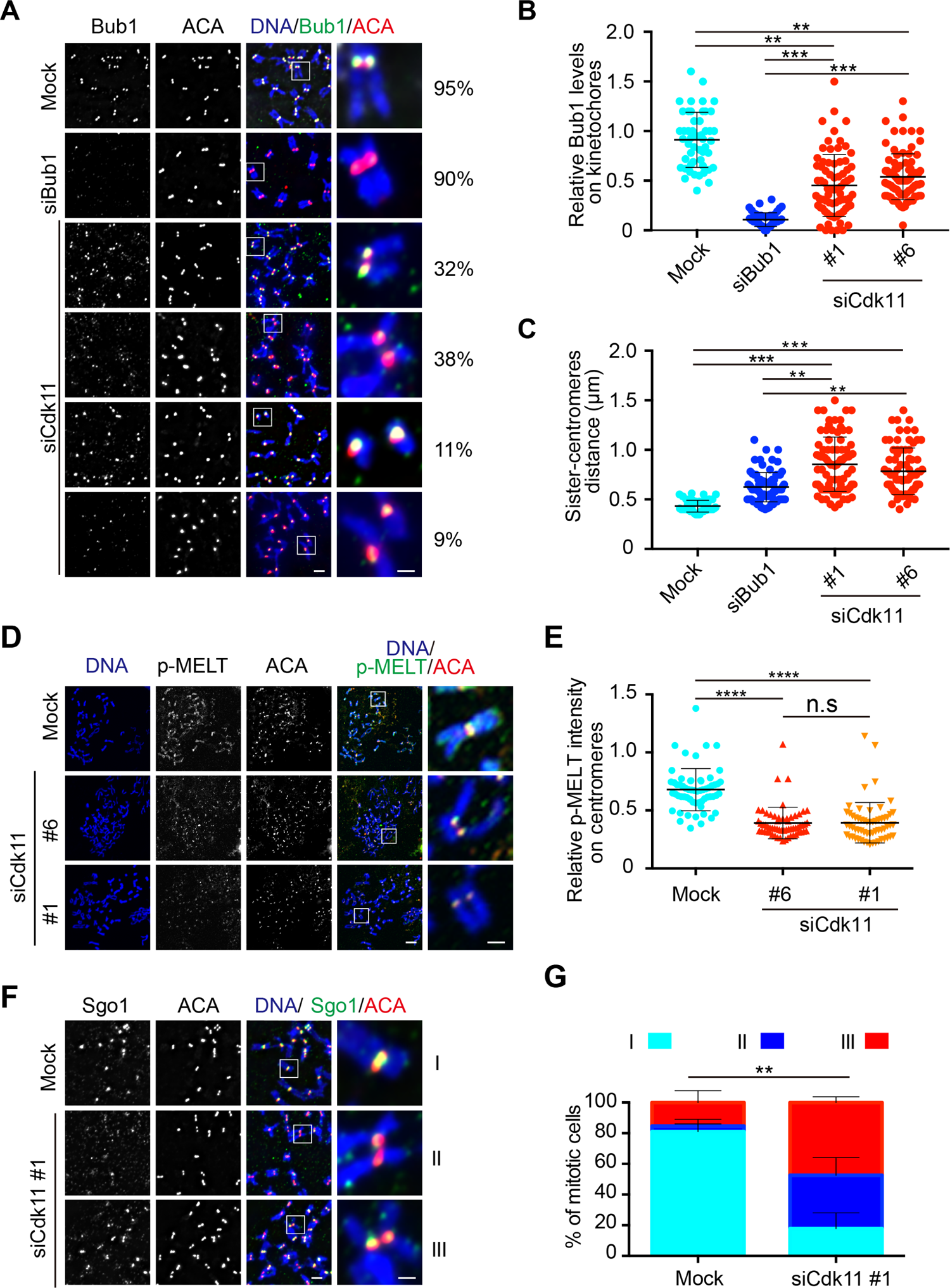
Cdk11 depletion partially decreases Bub1 and Knl1 p-MELT levels on kinetochores. **A.** Bub1 levels on kinetochores are marginally reduced by Cdk11 depletion. Nocodazole-arrested HeLa Tet-On cells were transfected with luciferase (mock), Bub1 or Cdk11 (#1 and #6) siRNAs and then subjected to chromosome spread and immunostaining with the indicated antibodies. Heterogeneity of Bub1 localizations on kinetochores in Cdk11 depletion cells was observed. Percentage of each type of Bub1 localization patterns is shown in the right panel: strong kinetochore Bub1 localization with weakened centromeric cohesion (32%), reduced kinetochore Bub1 localization with weakened centromeric cohesion (38%), strong kinetochore Bub1 localization with severely impaired centromeric cohesion (11%), and reduced kinetochore Bub1 localization with severely impaired centromeric cohesion (9%). Scale bars, 5 μm and 1 μm, respectively. **B** and **C.** Quantifications of relative Bub1 levels (Bub1/ACA, **B**) and the distance of inter-sister centromeres (**C**) in (**A**). These quantifications were performed based on a single experiment. At least 90 centromeres (6 per cell) were scored for each condition in (**B** and **C**). The average and standard deviation are shown here. The experiment was repeated three times and the results were highly reproducible. Quantification details hereafter were recorded in the section of Methods and Metatrails**. D.** Knl1 p-MELT levels on kinetochores are reduced by Cdk11 depletion. Nocodazole-arrested HeLa Tet-On cells were transfected with luciferase (mock) and Cdk11 (#1 and #6) siRNAs and then subjected to chromosome spread and immunostaining with the indicated antibodies. Scale bars, 5 μm and 1 μm, respectively. **E.** Quantification of relative p-MELT intensity (p-MELT/ACA) in (**D**). The quantification was performed based on a single experiment. At least 60 centromeres (5 per cell) were scored for each condition. The average and standard deviation are shown here. The experiment was repeated three times and the results were highly reproducible. **F.** Sgo1 localization in Cdk11 depletion cells. Nocodazole-arrested HeLa Tet-On cells were transfected with luciferase (mock) or Cdk11 (#1) siRNAs and then subjected to chromosome spread and immunostaining with the indicated antibodies. Scale bars, 5 μm and 1 μm, respectively. **G.** Quantification of Sgo1 localization patterns in Cdk11 depletion cells. Three types of Sgo1 localization patterns were observed: strong inner-centromeric Sgo1 localization with normal centromeric cohesion (type I), reduced centromeric Sgo1 localization with weakened centromeric cohesion (type II), and robust centromeric Sgo1 localization with weakened centromeric cohesion (type III). The quantification was carried out based on three independent repeats. The average and standard deviation calculated from at least three independent experiments are shown here. At least 200 sister-centromere pairs (10 per cell) were scored for each condition in a single repeat. n.s. denotes not significant; **, P<0.01; ***, P<0.001; ****, P<0.0001.

Bub1 enriches Sgo1 to centromeres during mitosis (Kawashima et al., 2010; Kitajima et al., 2005a; Tang et al., 2004a). If Bub1 decrease from kinetochores was responsible for Cdk11 depletion-induced centromeric cohesion defects, Sgo1 would also be expected to be dislodged from centromeres. We then examined Sgo1 localization in Cdk11 depletion cells. HeLa Tet-On cells were transfected with luciferase (mock) or Cdk11 siRNAs. Cells were then treated with nocodazole for 2 hr and mitotic cells were collected for chromosome spread and immunostaining. In more than 80% of nocodazole-arrested mock-treated HeLa Tet-On cells, Sgo1 localized to inner centromeres with robust centromeric cohesion (**Figures 1F** and **1G**). In contrast, three types of Sgo1 localization patterns were observed in nocodazole-arrested Cdk11 depletion cells: strong inner-centromeric Sgo1 localization with normal centromeric cohesion (type I, ∼20%), reduced centromeric Sgo1 localization with weakened centromeric cohesion (type II, ∼35%), and robust centromeric Sgo1 localization with weakened centromeric cohesion (type III, ∼45%). Again, there seemed to be no obvious correlation between centromeric Sgo1 levels and the robustness of centromeric cohesion in Cdk11 depletion cells. Thus, the Bub1-Sgo1 pathway is unlikely the major factor contributing to Cdk11-regulated centromeric cohesion.

### Ectopic targeting of Bub1 to kinetochores marginally rescues centromeric cohesion defects upon Cdk11 depletion

If Bub1 delocalization from kinetochores was the major factor contributing to Cdk11 depletion-induced centromeric cohesion defects, artificial restoration of Bub1 levels on kinetochores would rescue the defects. To test this, we ectopically targeted Bub1 to kinetochores by fusing Bub1 with Mis12, a kinetochore protein. We firstly determined the functionality of the fusion protein Mis12-Bub1 by examining to what extent it was able to rescue Bub1 depletion phenotypes. HeLa Tet-On cells depleted of Bub1 were transfected with plasmids containing GFP-Mis12-Bub1 (kinase domain, resides: 633-1085) WT or KD (kinase dead). Cells were then treated with nocodazole for 2 hr and mitotic cells were collected for chromosome spread followed by immunostaining and Sgo1 localization at centromeres was examined. Consistent with the previous findings (Kitajima et al., 2005b; Liu et al., 2013b; Tang et al., 2004b; Williams et al., 2017), Bub1 depletion dramatically decreased Sgo1 levels from centromeres (**Figures 2A, 2B** and **S1A**). Expression of GFP-Mis12-Bub1 WT, not KD, completely restored Sgo1 levels in Bub1-depleted cells (**Figures 2A, 2B** and **S1A**). GFP-Mis12-Bub1 WT and KD both localized to kinetochores similarly (**Figure 2B, low panel**). Thus, Mis12 is able to effectively target fully functional Bub1 to kinetochores. Then, we examined the extent to which these fusion proteins rescued the phenotypes of Cdk11 depletion. Consistently, Cdk11 depletion moderately decreased Sgo1 localization on centromeres and significantly weakened centromeric cohesion (**Figures 2C, 2D and S1C**). Expression of GFP-Mis12-Bub1 WT, not KD, fully restored centromeric Sgo1 levels but only partially rescued the centromeric cohesion defects (**Figures 2D, S1B** and **S1C**). Detailed analyses revealed that expression of WT only reduced the number of type II cells (from ∼20% to ∼5%), but barely on type III (from ∼45% to ∼48%) (**Figure 2E**). These results further confirm that the Bub1-Sgo1 pathway only partially contributes to Cdk11-regulated centromeric cohesion. Other unknown mechanisms are yet to be uncovered.

**Figure 2.**
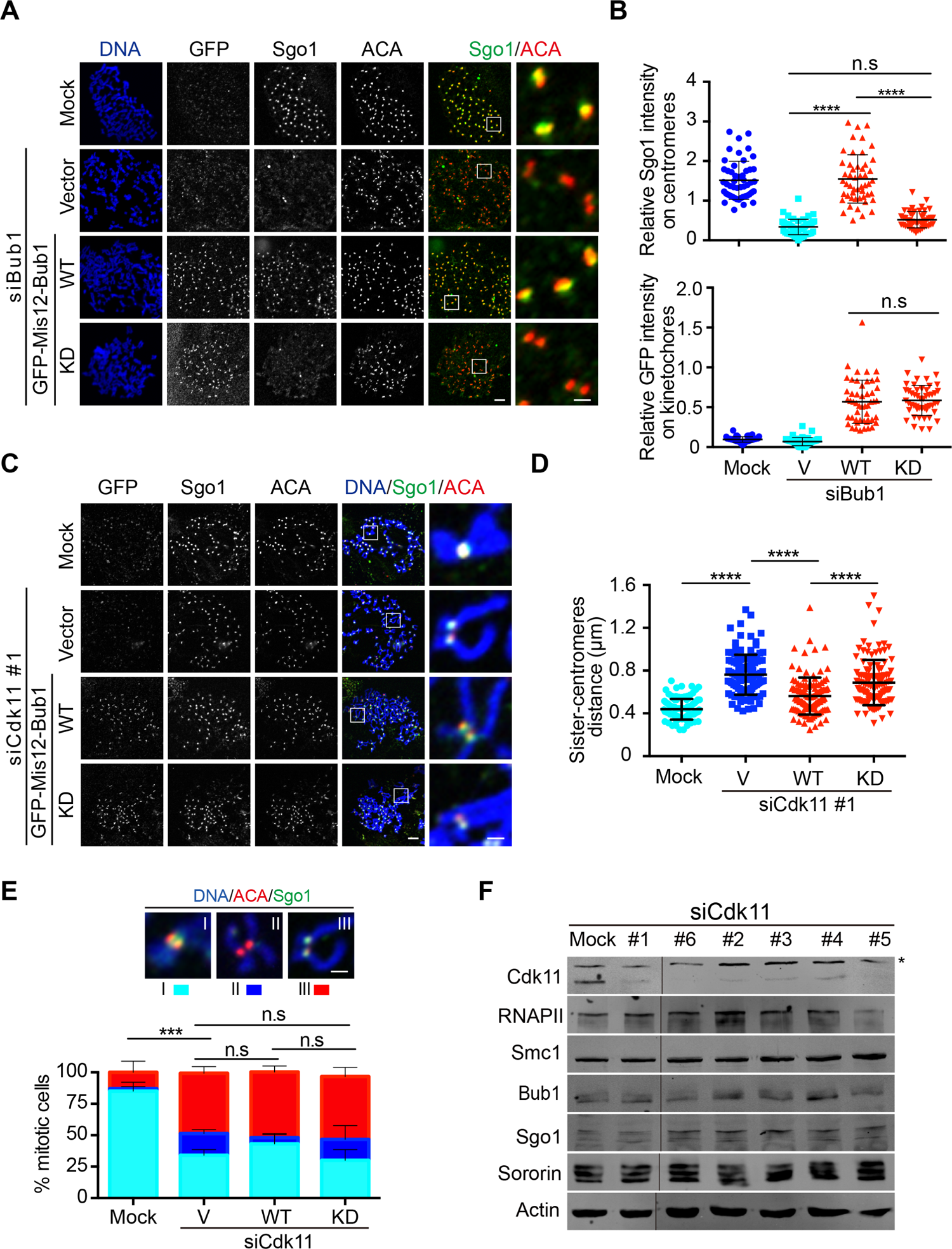
Ectopically targeting Bub1 to kinetochore slightly relieves centromeric cohesion defects in Cdk11 depletion cells. **A.** Targeting Bub1 kinase domain to kinetochores by Mis12 fully rescues Sgo1 localization defects on kinetochores in Bub1 depletion cells. HeLa Tet-On cells treated with luciferase (mock) or Bub1 siRNAs were transfected with vector or GFP-Mis12-Bub1 (631-1085) WT or KD (D946N). Cells were then treated with nocodazole for 2 hr and mitotic cells were collected for chromosome spread and immunostaining with the indicated antibodies. Scale bars, 5 μm and 1 μm, respectively. **B.** Quantifications of relative Sgo1 levels on centromeres (Sgo1/ACA, top panel) and GFP levels on kinetochores (GFP/ACA, bottom panel) in (**A**). These quantifications were performed based on a single experiment. The average and standard deviation are shown here. At least 50 centromeres (5 per cell) were scored for each condition. The experiment was repeated three times and the results were highly reproducible. **C.** Targeting Bub1 kinase domain to kinetochores by Mis12 slightly rescues centromeric cohesion defects in Cdk11 depletion cells. HeLa Tet-On cells treated with luciferase (mock) or Cdk11 siRNAs were transfected with vector or GFP-Mis12-Bub1 (631-1085) WT or KD. Cells were then treated with nocodazole for 2 hr and mitotic cells were collected for chromosome spread and immunostained with the indicated antibodies. Scale bars, 5 μm and 1 μm, respectively. **D.** Quantification of the distance of inter-sister centromeres in (**C**). The quantification was performed based on a single experiment. At least 100 centromeres (10 per cell) were scored for each condition. The average and standard deviation are shown here. The experiment was repeated three times and the results were highly reproducible. **E.** Quantification of chromosome morphology with distinct Sgo1 localization patterns (types I, II and III) in (**C**), described in (Figure 1F). The average and standard error calculated from three independent experiments are shown here. Scale bar, 1 μm. **F.** Lysates of HeLa Tet-On cells treated with mock or distinct Cdk11 siRNAs were resolved with SDS-PAGE and blotted with the indicated antibodies. Asterisk indicates non-specific protein bands. The black line indicates the membrane was cropped. The blots were run under the same experimental conditions and cropped from same membrane. n.s. denotes not significant; ***, P<0.001; ****, P<0.0001.

Cdk11 is an important regulator of mRNA splicing and processing (Dickinson et al., 2002; Hu et al., 2003; Loyer et al., 2008; Loyer et al., 1998; Trembley et al., 2002; Valente et al., 2009). Defective mRNA splicing has been shown to decrease the protein levels of an essential cohesion protector Sororin, leading to centromeric cohesion defects (Oka et al., 2014; Sundaramoorthy et al., 2014; van der Lelij et al., 2014; Watrin et al., 2014). We therefore examined if Cdk11 depletion-induced centromeric cohesion defects were a consequence of reduced protein levels of Sororin and/or other cohesion regulators. Western-blot analyses demonstrated that none of Smc1, Bub1, Sgo1, and Sororin protein levels were significantly changed in cells with Cdk11 depletion by several distinct siRNA oligos (**Figure 2F**). Thus, the protein levels of cohesin subunits and critical regulators are unlikely involved in Cdk11-regulated centromeric cohesion. However, we cannot completely exclude the possibility that Cdk11-mediated mRNA splicing of other factors contribute to the maintenance of centromeric cohesion.

### Cdk11 and its kinase activity maintain RNAPII and RNAPII-pSer2 on centromeres throughout the cell cycle

Centromeric transcription maintains centromeric cohesion and Cdk11 is a transcription regulator (Chen et al., 2021; Loyer and Trembley, 2020). We therefore hypothesized that Cdk11 promotes centromeric transcription to maintain centromeric cohesion. To test this hypothesis, we firstly examined how Cdk11 regulates RNAPII and elongating RNAPII (pSer2) as both of them are enriched at human centromeres during mitosis (Chan et al., 2012; Chen et al., 2021; Liu et al., 2015; Perea-Resa et al., 2020). Nocodazole-arrested HeLa Tet-On cells were transfected with luciferase (mock) or distinct Cdk11 siRNAs. Mitotic cells were subjected to chromosome spread and immunostaining. Two different RNAPII antibodies were used to recognize total (4H8) and phospho-Ser2 (H5) of RNAPII at centromeres (Chan et al., 2012; Chen et al., 2021; Liu et al., 2015). Consistently, robust RNAPII and RNAPII-pSer2 signals were detected in mock cells (**Figures 3A** and **3E**). Encouragingly, Cdk11 depletion by two distinct siRNAs largely decreased both RNAPII and RNAPII-pSer2 signals on centromeres (**Figures 3B** and **3F**); at the same time, these cells suffered centromeric cohesion defects (**Figure 3G**). Decreased elongating RNAPII-pSer2 levels and weakened centromeric cohesion were also observed in nocodazole-arrested mitotic non-transformed RPE1 cells depleted of Cdk11, suggesting that Cdk11 depletion-caused phenotypes are not cell type-specific (**Figures 3C, 3D** and **S2A**). In addition, using chromatin immunoprecipitation, we also found that Cdk11 depletion also decreased RNAPII-pSer2 levels on centromeres, but did not do so on two intergenetic regions in Log-phase HeLa Tet-On cells (**Figures 3H** and **S2D**). Thus, Cdk11 promotes centromeric transcription likely throughout the cell cycle.

**Figure 3.**
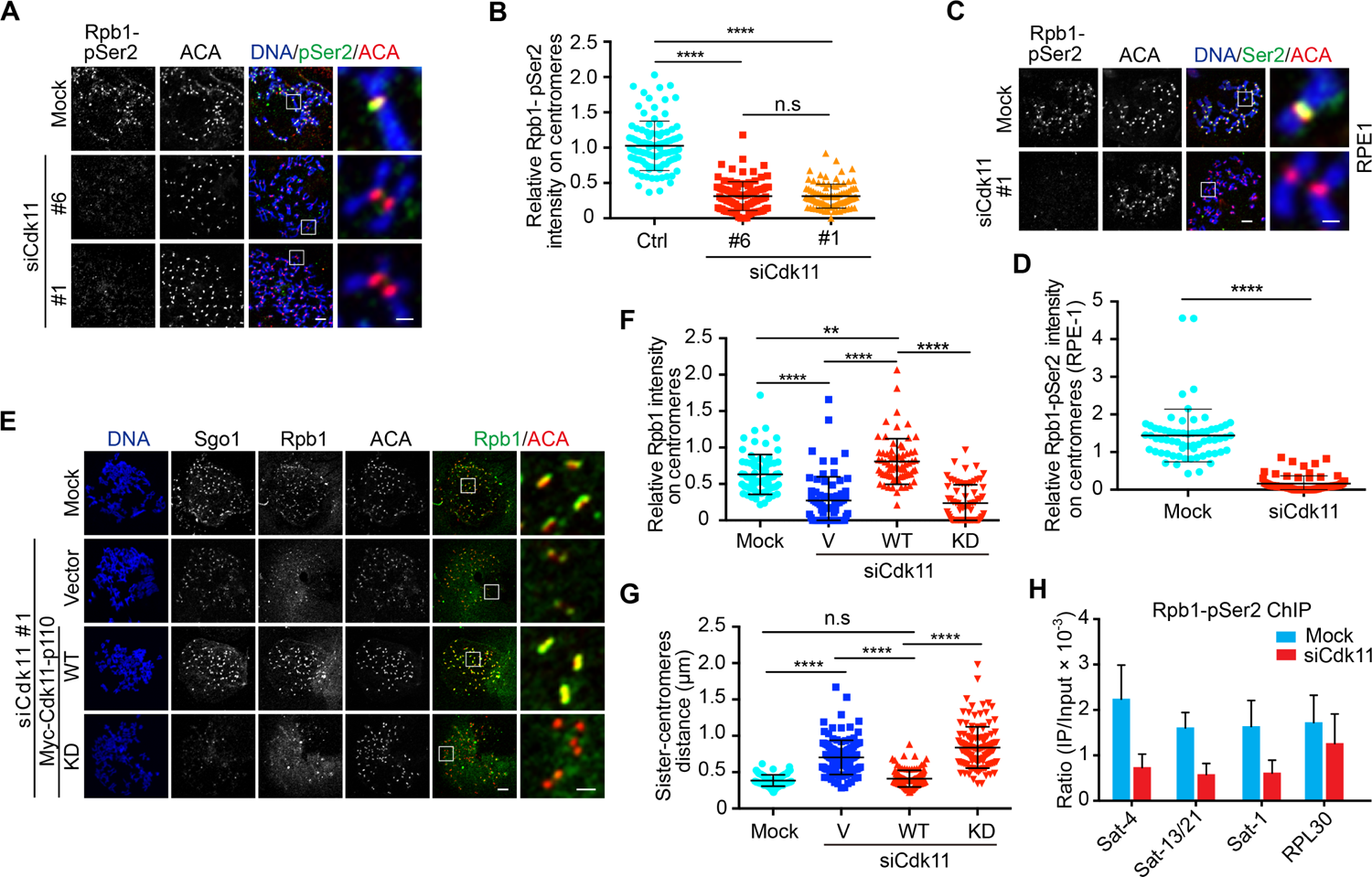
RNAPII and RNAPII pSer2 levels on centromeres are reduced by Cdk11 depletion. **A.** Cdk11 decreases RNAPII pSer2 (Rpb1-pSer2) levels on centromeres in HeLa Tet-On cells during mitosis. Nocodazole-arrested HeLa Tet-On cells were transfected with luciferase (mock) or distinct Cdk11 siRNAs. Mitotic cells were subjected to chromosome spread and immunostaining with the indicated antibodies. Antibody H5 was used to recognize RNAPII phosphorylated at Ser2. Scale bars, 5 μm and 1 μm, respectively. **B.** Quantification of relative Rpb1-pSer2 (Rpb1-pSer2/ACA) in (**A**). The quantification was performed based on a single experiment. At least 90 centromeres (6 per cell) were scored for each condition. The average and standard deviation are shown here. The experiment was repeated twice and the results were highly reproducible. **C** and **D.** Cdk11 decreases RNAPII pSer2 (Rpb1-pSer2) levels on centromeres in RPE-1 cells during mitosis. RPE-1 cells treated with the same condition as in (**A**) were subjected to chromosome spread and immunostaining with the indicated antibodies (**C**). Scale bars, 5 μm and 1 μm, respectively. Quantification of relative Rbp1-pSer2 is shown in (**D**). The quantification was performed based on a single experiment. At least 66 centromeres (6 per cell) were scored for each condition. The average and standard deviation are shown here. The experiment was repeated twice and the results were highly reproducible. **E.** Cdk11 activity maintains RNAPII (Rpb1) on centromeres in HeLa Tet-On cells during mitosis. HeLa Tet-On cells treated with luciferase (mock) or Cdk11 siRNAs were transfected with vector or Myc-Cdk11-p110 WT or KD. Cells were then treated with nocodazole for 2 hr and mitotic cells were collected for chromosome spread and immunostaining with the indicated antibodies. Antibody 4H8 was used to recognize total RNAPII. Scale bars, 5 μm and 1 μm, respectively. **F** and **G**. Quantifications of relative Rpb1 levels (Rpb1/ACA, **F**) and the distance of inter-sister centromeres (**G**) in (**E**). These quantifications were performed based on a single experiment. At least 60 centromeres (5 per cell) for each condition in (**F**) and at least 120 sister-centromere pairs (10 per cell) for each condition in (**G**) were scored. The average and standard deviation are shown here. The experiment was repeated three times and the results were highly reproducible**. H.** RNAPII pSer2 is associated with centromeric chromatin. Log-phase HeLa Tet-On cells were crosslinked with formaldehyde and the subsequent lysates were subjected to chromatin immunoprecipitation (ChIP) assay. Antibody Active Motif was used to recognize RNAPII phosphorylated at Ser2. Immunoprecipitated (IP) DNA fragments were analyzed with the indicated primers by real-time PCR. The average of ratios (IP/Input × 10^-3^) and stand error calculated from three independent experiments are shown here. n.s. denotes not significant; **, P<0.01; ****, P<0.0001.

We next determined whether Cdk11 kinase activity is required for RNAPII localization on centromeres, HeLa Tet-On cells stably expressing Myc-Cdk11-WT, or kinase dead mutant (KD) were depleted of endogenous Cdk11 and then incubated with nocodazole for 2 hr before harvest for chromosome spread and immunostaining. Western-blot analyses showed that Myc-Cdk11 WT and KD were expressed at the comparable level in Cdk11-depleted cells (**Figure S2B**). Expression of Myc-Cdk11 WT completely restored RNAPII levels on centromeres, but Cdk11 KD failed to do so (**Figures 3E** and **3F**), suggesting that the Cdk11 kinase activity is required for centromeric localization of RNAPII. Accordingly, expression of Cdk11 WT, not KD, also completely rescued centromeric cohesion defects (**Figures 3G**). Cdk11-p58 isoform is specifically expressed at G2/M phase and plays an essential role in protecting centromeric cohesion during mitosis (Hu et al., 2007; Rakkaa et al., 2014a). Then we examined whether the kinase activity of Cdk11-p58 was also required for maintaining RNAPII at mitotic centromeres. Cdk11-depleted Hela Tet-on cells were transfected with vector, myc-Cdk11-p58 WT or kinase dead mutant (KD) and incubated with nocodazole for 2 hr before harvest for chromosome spread and immunostaining. Similarly, Cdk11-p58 WT completely restored RNAPII levels and centromeric cohesion defects in Cdk11 depletion cells but KD failed to do so (**Figures S3A**-**3D**). Altogether, we conclude that Cdk11 and its activity are required for maintaining active RNAPII on centromeres throughout the cell cycle, and that Cdk11-promoted centromeric transcription is required for maintaining centromeric cohesion.

### Mitosis-specific degradation of Cdk11 result in centromeric cohesion defects and dislodge RNAPII from centromeres

As Cdk11 is also required for maintaining RNAPII on centromeres in interphase, reduced RNAPII levels on centromeres in mitosis could be a legacy inherited from interphase. Therefore, we sought to determine whether mitotic Cdk11 is required for maintaining RNAPII on centromeres and also to examine centromeric cohesion during mitosis by mitosis-specifically degrading G2/M Cdk11-p58. HeLa Tet-On cells stably expressing the auxin (indole-3 acid acid, IAA) receptor Myc-TiR1 were depleted of endogenous Cdk11 and transfected with GFP-AID-Cdk11-p58. Nocodazole-enriched mitotic cells were collected and treated with IAA for 3 hr (**Figure 4A**). Western-blot analyses demonstrated that GFP-AID-Cdk11-p58 protein levels were largely degraded 3 hr after auxin treatment (**Figure 4B**), validating this system. Consistently, expression of GFP-AID-Cdk11-p58 completely rescued the reduced RNAPII levels on centromeres and centromeric cohesion defects in Cdk11-depleted cells (**Figures 4C, 4D** and **4E**). Strikingly, degradation of GFP-AID-Cdk11-p58 by IAA treatment recapitulated the defects by Cdk11, thus supporting that mitotic Cdk11-p58 is required for maintaining centromeric RNAPII and centromeric cohesion during mitosis.

**Figure 4.**
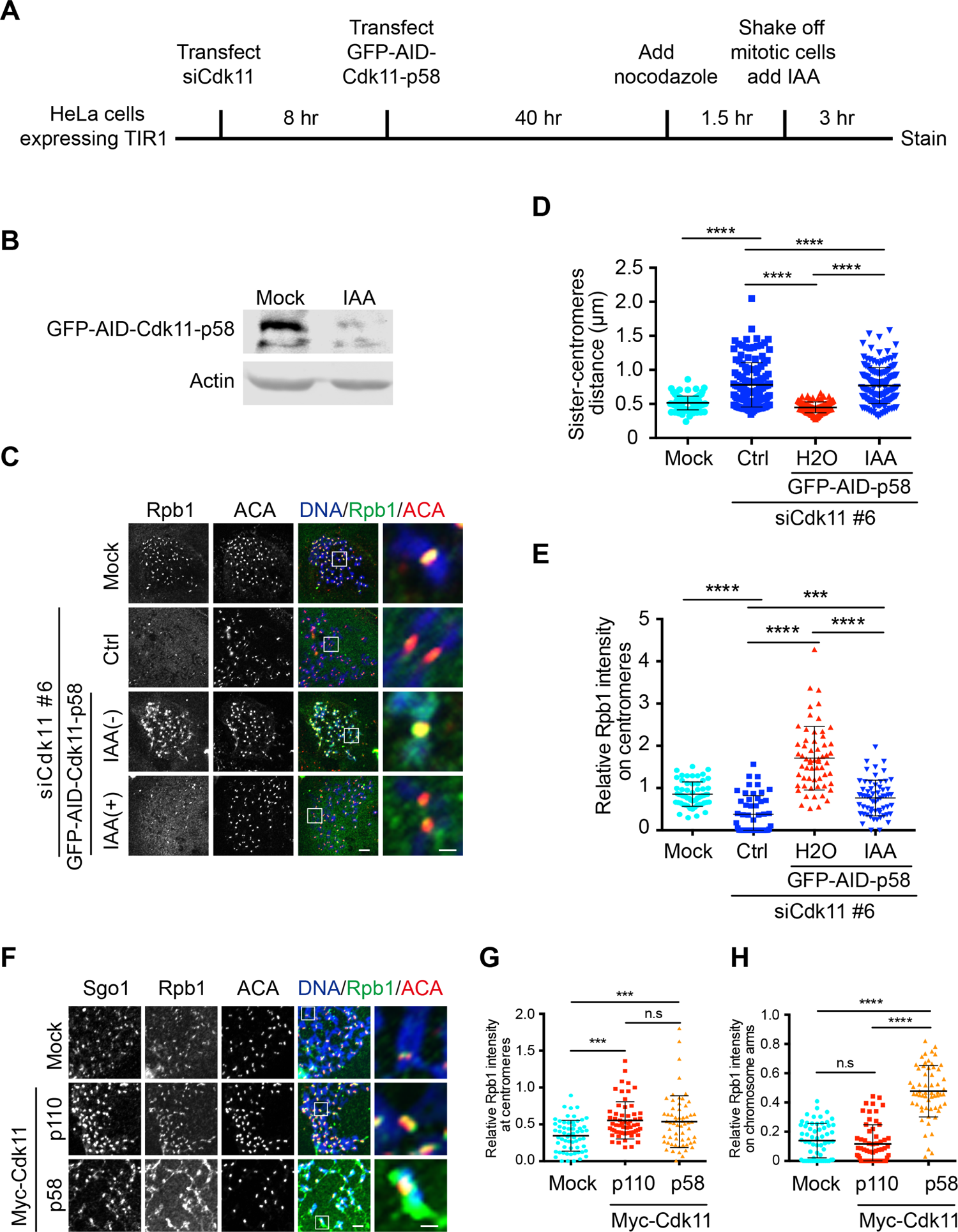
Mitosis-specific degradation of Cdk11-p58 or inhibition of Cdk11 reduces RNAPII levels on centromeres and weakens centromeric cohesion. **A.** Schematic of Auxin (IAA)-inducible degradation of Cdk11-p58 in mitosis. **B** and **C.** IAA-induced degradation of Cdk11-p58 reduces RNAPII levels on centromeres and weakens centromeric cohesion during mitosis. HeLa Tet-On cells stably expressing Myc-Tir1 were treated with luciferase (mock) or Cdk11 siRNAs and transfected with vectors or plasmids containing GFP-AID-Cdk11-p58. Nocodazole-arrested mitotic cells were collected and further treated with IAA for 3 hr. Cells were finally subjected to immunostaining with the indicated antibodies in (**C**) or analyzed by Western blotting (**B**). Scale bars, 5 μm and 1 μm, respectively. **D** and **E.** Quantifications of relative Rpb1 levels (Rpb1/ACA, **E**) and the distance of inter-sister centromeres (**D**) in (**C**). These quantifications were performed based on a single experiment. At least 110 sister-centromere pairs (10 per cell) for each condition in (**D**) and at least 60 centromeres (5 per cell) for each condition in (**E**) were scored. The average and standard deviation are shown here. The experiment was repeated three times and the results were highly reproducible. **F.** Cdk11-p58 overexpression increases RNAPII levels on both centromeres and chromosome arms. Inducible HeLa cells Tet-On cells with Myc-Cdk11-p110 or Myc-Cdk11-p58 were treated with doxycycline. Cells were treated with nocodazole and then subjected to chromosome spread and immunostaining with the indicated antibodies. Scale bars, 5 μm and 1 μm, respectively. **G** and **H**. Quantification of relative Rpb1 levels on centromeres (Rpb1/ACA, **G**) and on chromosome arms (Rpb1/DNA, **H**). These quantifications were performed based on a single experiment. At least 60 centromeres (5 per cell, **G**) and 60 chromosome arms (5 per cell, **H**) were scored for each condition. The average and standard deviation are shown here. The experiment was repeated twice and the results were highly reproducible. n.s. denotes not significant; **, P<0.01; ***<0.001 ****, P<0.0001.

As Cdk11 depletion decreased centromeric RNAPII levels in mitosis, we reasoned that Cdk11 overexpression could increase them. To test this, we examined centromeric RNAPII in nocodazole-arrested HeLa Tet-On cells stably expressing Myc-Cdk11-p110 and p58. As expected, expression of both isoforms of Cdk11 augmented RNAPII signals at centromeres (**Figures 4F, 4G** and **S4**). Interestingly, ectopic RNAPII was also observed on chromosome arms in some of Cdk11-p58 cells (**Figure 4H**).

### Cdk11 physically associates with RNAPII via its C-terminus *in vivo* and is present at centromeric chromatin

Cdk11 was shown to bind the CTD of RNAPII *in vitro* (Trembley et al., 2003), and phosphorylate RNAPII at Ser2 (Gajduskova et al., 2020). We then examined how Cdk11 physically interacted with RNAPII in cells. HeLa Tet-On cells stably expressing Myc-Cdk11-p110 WT or KD were cross-linked with formaldehyde and the resulting cell lysates were subjected to RNAPII immunoprecipitation. Consistent with *in vitro* results, both Myc-Cdk11-p110 WT and KD bound with RNAPII (**Figure 5A**). This result also suggests that the Cdk11-RNAPII binding is independent of Cdk11 kinase activity. In addition, Cdk11-p58 also physically interacted with RNAPII (**Figure 5B**). As Cdk11-p58 shares the same C-terminus with Cdk11, it is very likely that Cdk11 binds to RNAPII through its C-terminus.

**Figure 5.**
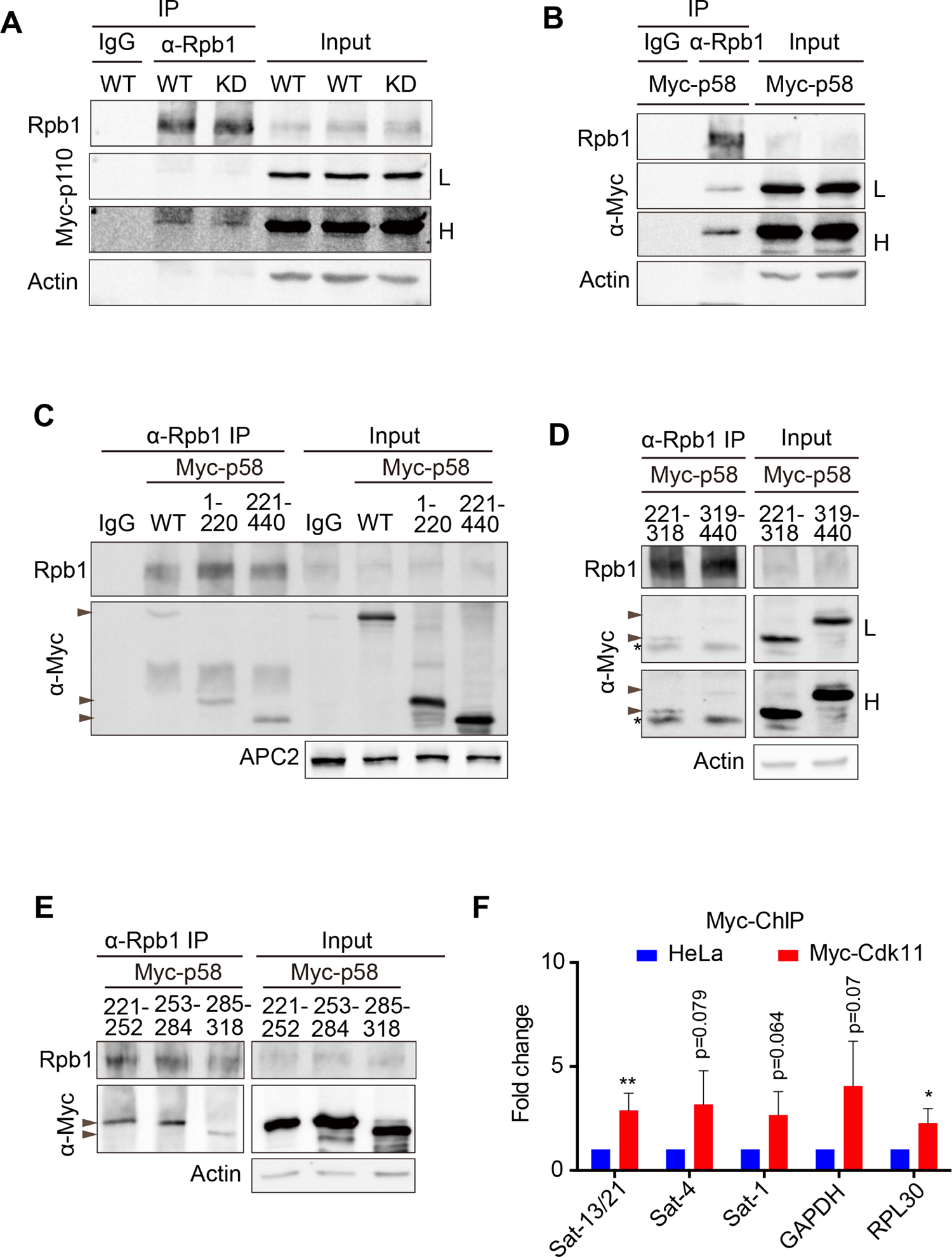
Cdk11 binds RNAPII and is associated with centromeric chromatin. **A** and **B.** Cdk11-p110 and Cdk11-p58 bind RNAPII. Log-phase inducible HeLa Tet-On cells with Myc-Cdk11-p110 (**A**) or p58 (**B**) were treated with doxycycline. The subsequent cell lysates were treated with IgG or antibody against Rpb1. Immunoprecipitated proteins were finally resolved with SDS-PAGE and blotted with indicated antibodies. L indicates light exposure and H indicates high exposure. **C.** Both N-terminal and C-terminal fragments of Cdk11-p58 bind RNAPII. Lysates of log-phase HeLa Tet-On cells transfected with plasmids containing Myc-Cdk11-p58, p58 (1-220) or p58 (221-440) were treated with IgG or antibody against Rpb1. The resulting immunoprecipitated proteins were resolved with SDS-PAGE and blotted with indicated antibodies. Arrowheads indicate the bindings. **D** and **E.** Further Mapping the domains in Cdk11-p58 binding to RNAPII. Lysates of log-phase HeLa Tet-On cells transfected with plasmids containing Myc-Cdk11-p58 (221-318) (**D**), p58 (319-440) (**D**), p58 (221-252) (**E**), p58 (253-284) (**E**), or p58 (285-318) (**E**), were treated with IgG or antibody against Rpb1. The resulting immunoprecipitated proteins were resolved with SDS-PAGE and blotted with indicated antibodies. L indicates light exposure and H indicates high exposure. Arrowheads indicate the bindings. Asterisk indicates the light chain of IgG. **F.** Cdk11 is associated with centromeric chromatin. Log-phase inducible HeLa Tet-On cells with Myc-Cdk11-p110 were treated with doxycycline. Collected cells were crosslinked with formaldehyde and the subsequent lysates were subjected to chromatin immunoprecipitation (ChIP) assay. Immunoprecipitated (IP) DNA fragments were analyzed with the indicated primers by real-time PCR. The average of normalized fold changes (IP/Input) and stand error calculated from three independent experiments are shown here. n.s. denotes not significant; *, P<0.05; **, P<0.01; ****, P<0.0001.

We then used Cdk11-p58 to further map the regions in Cdk11 responsible for RNAPII binding. Myc-Cdk11-p58 N-terminal (1-220) and C-terminal (221-440) fragments were constructed and their interactions with RNAPII were tested. Both fragments were immunoprecipitated by RNAPII, but the C-terminus (residues 221-440) showed a slightly stronger association with RNAPII (**Figure 5C**). With further mapping, we were finally able to identify two regions of Cdk11-p58 containing residues 221-252 and 253-284 that are responsible for RNAPII binding (**Figures 5D** and **5E**). Notably, these two regions are highly conserved across species and contain Cdk11 kinase domain as well (**Figure S5A**), suggesting that the Cdk11-RNAPII binding is likely conserved.

We next determined if Cdk11 could localize to centromeres as it regulates centromeric RNAPII. Immunostaining was performed to examine the localization of Myc-Cdk11-p58 in HeLa Tet-On cells. In interphase cells, Myc-Cdk11 p58 localized in both cytoplasm and nucleus; in mitosis, it was mainly enriched at the edges surrounding chromosomes but was undetectable on chromosomes (**Figure S5B**). We then performed chromatin-immunoprecipitation (ChIP) to detect the presence of Cdk11 on centromeres. As shown in **Figure 5F**, Myc-Cdk11 was detected at both the tested gene regions and centromeres, suggesting that Cdk11 can localize to centromeres albeit its localization may not be specific to centromeres. Low abundance on chromosomes might explain the failure in its detection on chromosomes by fluorescence.

### Cdk11 depletion and expression of its kinase-dead version decreases centromeric alpha-satellite RNAs

As Cdk11 is required for maintaining active RNAPII on centromeres, we then sought to determine how Cdk11 regulates centromeric RNA transcripts. Log-phase HeLa Tet-on cells were depleted of Cdk11 and total RNAs were extracted for real-time PCR analysis using two gene primers (GAPDH and RPL30) and two centromeric alpha-satellite primers (Sat-4 and Sat-13/21). Cdk11 depletion with four distinct siRNA oligos had minimal effects on the amounts of gene RNAs, but unanimously reduced the amounts of centromeric RNAs on Sat-4 and Sat-13/21 to varying levels (**Figure 6A**), suggesting that Cdk11 promotes centromeric transcription. To further determine if Cdk11 is required for transcriptional activity on centromeres, we examined 5’-Ethynyl Uridine (EU)-labelled nascent RNA transcripts. Hela Tet-on cells were depleted of Cdk11 and chased with EU 1 hr before harvest. EU-labelled RNAs were then purified and subjected to real-time PCR analysis. Cdk11 depletion slightly decreased the amount of EU-RNAs on GAPDH and barely affected that of RPL30 EU-RNAs (**Figure 6B**). In contrast, Cdk11 depletion significantly decreased the amounts of the two tested centromeric alpha-satellite EU-RNAs. These results, together with its requirement in maintaining active RNAPII on centromeres, indicate that Cdk11 promotes centromeric transcription directly via RNAPII.

**Figure 6.**
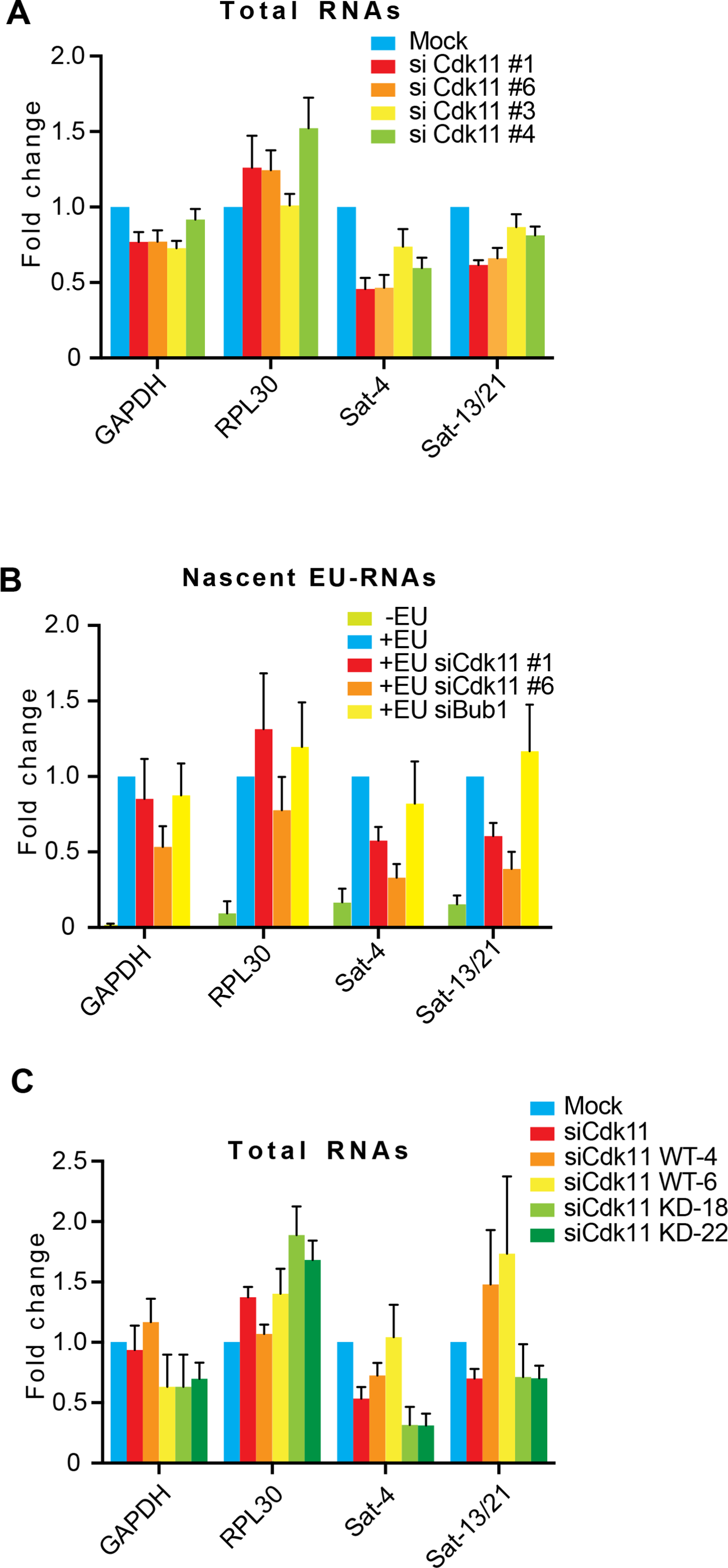
Cdk11 depletion or inhibition decreases centromeric alpha-satellite RNAs. **A.** Cdk11 depletion reduces the amount of centromeric alpha-satellite RNAs. Total RNAs were extracted from HeLa Tet-On cells transfected with mock or distinct Cdk11 siRNAs. Real-time PCR was performed to evaluate the amount of RNAs as indicated. The average and stand error calculated from four independent experiments are shown here. **B.** Cdk11 depletion decreases the amount of centromeric alpha-satellite nascent EU-RNAs. Log HeLa Tet-On cells transfected with luciferase (mock) or distinct Cdk11 siRNAs were treated with EU 1 hr before harvest. EU-RNAs were purified and then subjected to real-time analysis with the indicated primers. The average and stand error calculated from at least four independent experiments are shown here. **C.** Cdk11 kinase activity is required for maintaining the amount of centromeric alpha-satellite RNAs. Log-phase inducible HeLa Tet-On cells with Myc-Cdk11-p110 WT (4 or 6) or KD (18 or 22) were treated with doxycycline and transfected with luciferase (mock) or Cdk11 siRNAs. Total RNAs were extracted from HeLa Tet-On cells transfected with mock or distinct Cdk11 siRNAs. Real-time PCR was performed to evaluate the amount of RNAs as indicated. The average and stand error calculated from at least three independent experiments are shown here.

To determine whether Cdk11 kinase activity is required for centromeric transcription, HeLa Tet-on cells stably expressing Myc-Cdk11 WT or KD were depleted of endogenous Cdk11, and total RNAs were extracted for real-time PCR analysis. Expression of Myc-Cdk11 WT, not KD, restored the decreased amounts of Sat-4 and Sat13/21 RNAs in Cdk11-depleted cells, while barely altered the expression of the tested genes (**Figure 6C**). Thus, Cdk11 and its kinase activity are required for efficient centromeric transcription.

Nucleoli were recently shown to be involved in the regulation of centromeric RNA levels (Bury et al., 2020). We then examined if Cdk11 depletion-reduced centromeric alpha-satellite RNAs could be a result of altered nucleoli. Cdk11 depletion seemed to have no impact on the morphology of nucleoli and the localization of a specific nucleolar protein, nucleolin (**Figure S5C**). Thus, nucleoli may not be involved in Cdk11-regulated centromeric transcription.

### Enhanced centromeric transcription rescues Cdk11 depletion defects

If Cdk11 depletion-induced centromeric cohesion defects were a consequence of decreased centromeric transcription, enhancing centromeric transcription should then be able to rescue the centromeric cohesion defects. We recently showed that THZ1 (Cdk7 inhibitor) treatment and expression of CENP-B DNA-binding domain (CENP-B DB) are two effective approaches to specifically enhance centromeric transcription (Chen et al., 2021). Therefore, we examined centromeric transcription and centromeric cohesion in Cdk11 depletion cells treated with THZ1 or expressing CENP-B DB. Consistently, Cdk11 depletion decreased the RNAPII levels on centromeres and the amounts of Sat-4 and Sat-13/21 RNAs (**Figures 7A, 7B, 7C** and **7E**). Interestingly, decreased centromeric transcription were both restored by THZ1 treatment or CENP-B DB expression and reduced RNAPII levels on centromeres were also restored by THZ1 treatment (**Figures 7A, 7B, 7C, 7E** and **S6**). At the same time, centromeric cohesion defects in Cdk11 depletion cells were also completely rescued (**Figures 7D** and **7F**). These strong genetic data further support that Cdk11 maintains centromeric cohesion mainly through promoting centromeric transcription (**Figure 7G**).

**Figure 7.**
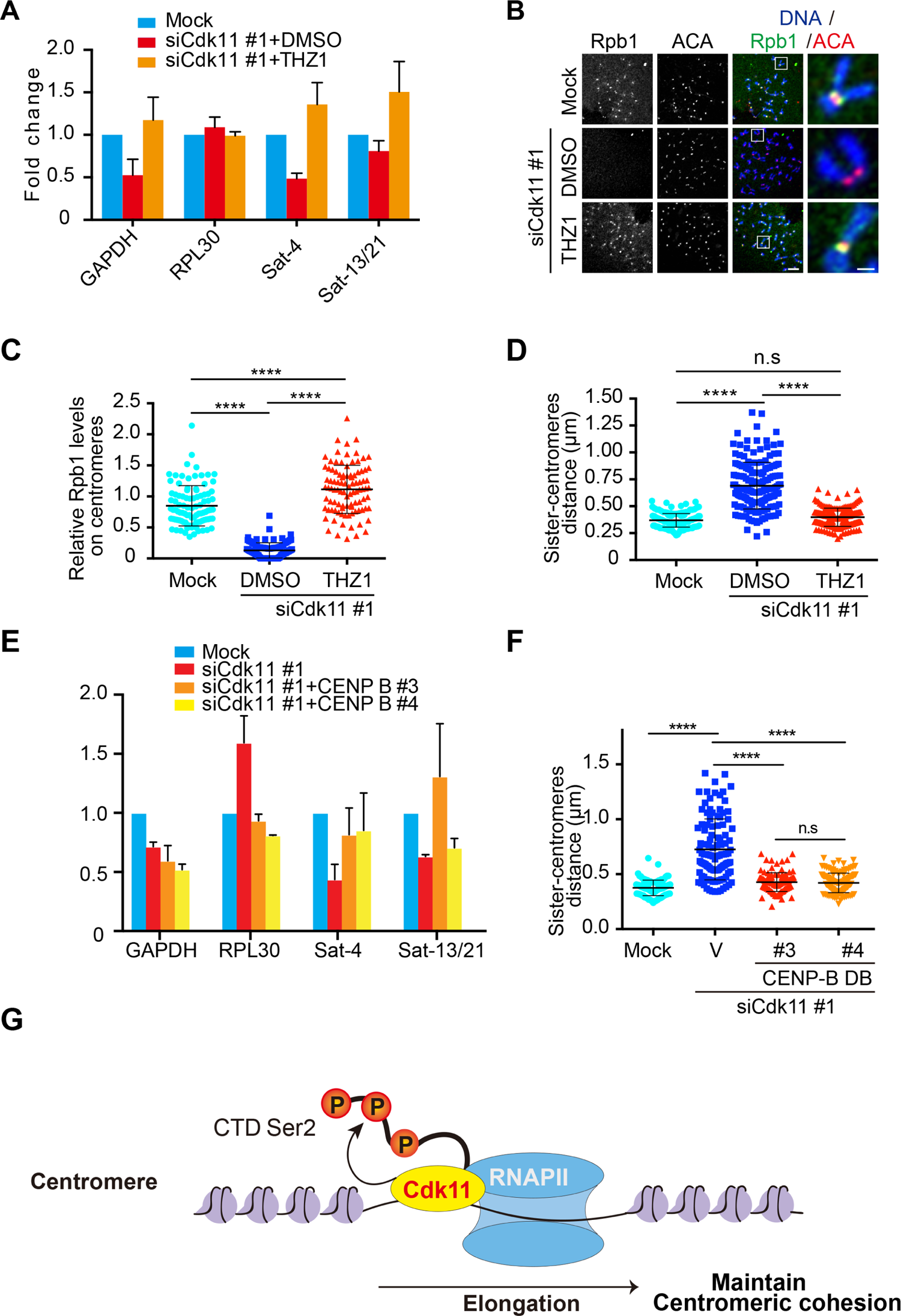
Ectopic expression of CENP-B DB or treatment with THZ1 rescues Cdk11 depletion-induced phenotypes. **A.** Treatment of THZ1 enhances centromeric transcription. Log-phase HeLa Tet-On cells transfected with luciferase (mock) or Cdk11 siRNAs (#1) were treated with DMSO or THZ1 for 12 hr and total RNAs were extracted for real-time PCR analysis. The average and stand error calculated from four independent experiments are shown here. **B.** Treatment of THZ1 rescues RNAPII localization defects and centromeric cohesion defects in Cdk11 depletion cells. HeLa Tet-On cells were arrested with thymidine and then released into fresh medium containing nocodazole. Cells were also treated with either DMSO or THZ1 12 hr before harvest. Collected mitotic cells were subjected to chromosome spread and immunostaining with the indicated antibodies. Scale bars, 5 μm and 1 μm, respectively. **C** and **D.** Quantifications of relative Rpb1 levels (Rpb1/ACA, **C**) and the distance of inter-sister centromeres (**D**) in (**B**). These quantifications were performed based on a single experiment. At least 95 centromeres (5 per cell) were scored for each condition in (**C**) and at least 167 sister-centromere pairs (10 per cell) were scored for each condition in (**D**). The average and standard deviation are shown here. The experiment was repeated twice and the results were highly reproducible. **E.** Expression of CENP-B DB rescues centromeric cohesion defects in Cdk11 depletion cells. Log-phase inducible HeLa Tet-On cells with Myc-CENP-B DB (#3 or #4) were treated with doxycycline and transfected with luciferase (mock) or Cdk11 siRNAs. Total RNAs were extracted from HeLa Tet-On cells transfected with mock or distinct Cdk11 siRNAs. Real-time PCR was performed to evaluate RNAs as indicated. The average and stand error calculated from three independent experiments are shown here. **F.** Quantification of the distance of inter-sister centromeres in CENP-B stable cells depleted of Cdk11. HeLa Tet-On cells with Myc-CENP-B DB were treated with doxycycline and were arrested with thymidine and then released into fresh medium containing nocodazole. Collected mitotic cells were subjected to chromosome spread and immunostaining with DAPI and ACA. The quantification was performed based on a single experiment. At least 110 centromeres (10 per cell) were scored for each condition. The average and standard deviation are shown here. The experiment was repeated twice and the results were highly reproducible. **G.** Working model. Constitutively expressed Cdk11 binds RNAPII and phosphorylates CTD-Ser2 at centromeres to promote centromeric transcription in both interphase and mitosis. This role of Cdk11 is redundant of Cdk9. G2/M-expressed Cdk11-p58 may facilitate the retention of RNAPII on centromeres during mitosis, thus helping maintain centromeric transcription and centromeric cohesion.

## Discussion

Centromeric transcription is a conserved process across species and plays an essential role in centromere functions, but a little is known about how centromeric transcription is regulated. We here identify Cdk11 as an important regulator for centromeric transcription. Cdk11 localizes to centromeres and interacts with and phosphorylates centromeric RNAPII to promote centromeric transcription, thus maintaining centromeric cohesion that is essential for faithful chromosome segregation in mitosis.

Ser2 phosphorylation of the RNAPII CTD is essential for RNAPII elongation during transcription. Cdk9 is the major kinase that phosphorylates that site, thus rendering Cdk9 a general transcriptional factor (Chou et al., 2020). As a relative to Cdk9, Cdk11 has also been shown to regulate transcription. However, distinct from Cdk9, Cdk11 appears not to universally regulate transcription; instead, it may do so only for a subset of genes (Drogat et al., 2012; Gajduskova et al., 2020). Consistent with this idea, our findings here suggest that Cdk11 is also required for the efficient transcription on a specialized chromosomal region, the centromere, thus establishing a novel role of Cdk11 in transcriptional regulation of non-coding DNA sequences. Mechanistically, Cdk11 binds and phosphorylates the RNAPII CTD to maintain elongating RNAPII on centromeres. However, it is worth mentioning that Cdk11-regulated transcription is not specific to centromeres and it is also important for the transcription of a subset of genes. In future, it would be of interest to understand how Cdk11 displays this type of specificity. Recently, we showed that Cdk9 is also critical for centromeric transcription (Chen et al., 2021). Thus, Cdk11 and Cdk9 may play a redundant role in promoting RNAPII elongation at centromeres. In support of this idea, Cdk11 depletion only resulted in a moderate reduction in centromeric transcription (**Figure 6**).

When cells enter mitosis, the majority of RNAPII and transcriptional factors are released from chromosomes, leading to a significant loss of RNAPII transcription on chromosomes (Palozola et al., 2017; Parsons and Spencer, 1997; Teves et al., 2018). However, robust levels of elongating RNAPII on centromeres are still remained and centromeres are under active transcription during mitosis (Bobkov et al., 2018; Chan et al., 2012; Chen et al., 2021; Liu et al., 2015; Perea-Resa et al., 2020). Thus, a mitosis-specific mechanism might exist to preserve RNAPII on centromeres, thus maintaining a considerable transcriptional activity. Cdk11 may be part of such a mechanism. In support of it, mitosis-specific degradation of G2/M Cdk11-p58 largely decreased the RNAPII levels at centromeres, and overexpression of Cdk11-p58 increased RNAPII levels on both centromeres and chromosome arms during mitosis (**Figure 4**). Of note, because our results indicate that Cdk11 activity is required for centromeric transcription likely throughout the cell cycle, it is possible that both Cdk11-p110 and -p58 are needed to maintain centromeric transcription during mitosis. Nevertheless, the isoform of p58 that is exclusively expressed in G2/mitosis may provide an extra insurance for the maintenance of centromeric transcription during mitosis.

How is Cdk11 involved in centromeric cohesion regulation? This question had not been satisfactorily addressed since the first discovery of Cdk11 involvement in the regulation of centromeric cohesion a decade ago. Although previous studies had thrown Bub1 under a spotlight (Hu et al., 2007; Rakkaa et al., 2014b), our data here suggest that Bub1 is not the answer. Specifically, ectopic restoration of fully functional Bub1 on kinetochores only marginally rescued centromeric cohesion defects in Cdk11 depletion cells, providing strong evidence against the main role of Bub1 in this regulation. Instead, we have here established Cdk11 as an important regulator for centromeric transcription and this role may empower Cdk11 to regulate centromeric cohesion. This conclusion is further supported by our genetic data showing that enhanced centromeric transcription completely rescued the weakened centromeric cohesion induced by Cdk11 depletion (**Figure 7**). Notably, previous studies showed that defective mRNA splicing induced centromeric cohesion defects through decreasing the protein levels of an essential cohesion protector Sororin (Oka et al., 2014; Sundaramoorthy et al., 2014; van der Lelij et al., 2014; Watrin et al., 2014). Considering a role of Cdk11 in mRNA splicing (Dickinson et al., 2002; Hu et al., 2003; Loyer et al., 2008; Loyer et al., 1998; Pak et al., 2015; Trembley et al., 2002; Valente et al., 2009), centromeric cohesion defects caused by Cdk11 could also be attributed to compromised mRNA splicing. Although our results here have largely excluded the involvement of Sororin and several other cohesion-regulators, we cannot completely rule out the possibility that Cdk11 promotes centromeric cohesion through regulating mRNA splicing for some yet-to-be-identified cohesion regulators. Nevertheless, our findings in this report provide a mechanism to explain why Cdk11 is important for centromeric cohesion and further highlight the emerging critical role of centromeric transcription in centromeric cohesion regulation (Chen et al., 2021; Liu, 2016; Liu et al., 2015). Notably, The centromeric transcription-cohesion relationship may also explain why a transcriptional regulator prohibitin 2 (PHB2) is important for centromeric cohesion regulation (Takata et al., 2007). In further, it would be of importance to identify more key factors that are specific and important for centromeric transcription as well as to explore the molecular mechanism through which centromeric transcription promotes centromeric cohesion.

## Experimental procedures

### Mammalian cell culture, siRNAs, and transfection

Hela Tet-on cells were cultured in Dulbecco’s modified Eagle’s medium (DMEM, Invitrogen) containing 10% fetal bovine serum and 10mM L-glutamine at 37°C and 5% CO_2_. RPE-1 cells were cultured in DMEM: F-12 medium (Invitrogen) supplemented with 10% FBS and 10 mM L-glutamine. To arrest cells at G1/S, cells were usually incubated in medium containing 2 mM thymidine (Sigma) for at least 16 h. Nocodazole was purchased from Sigma Aldrich. OTS964 was purchased from MedChemExpress. THZ1 was purchased from Selleckchem.

Plasmid transfection was implemented using the Effectene reagent (Qiagen) according to the manufactures’ protocols. For Myc-Cdk11 (WT or KD) or Myc-Cdk11-p58 WT stable cells, Hela Tet-on cells were transfected with pTRE2 vectors encoding RNAi-resistant Myc-Cdk11 or Myc-Cdk11-p58 and selected with 350 μg ml^−1^ hygromycin (Invitrogen). The surviving clones were screened for expression of the desired proteins in the presence of 1 μg ml^−1^ doxycycline (Invitrogen). Expression of Myc-Cdk11 was also induced with 1 μg ml^−1^ doxycycline in the subsequent experiments.

For RNAi experiments, siRNA oligonucleotides were purchased from Dharmacon. HeLa or RPE-1 cells were transfected using Lipofectamine RNAiMax (Invitrogen) according to the manufacturer’s protocols. Subsequent analyses were usually performed 48 hr after transfection with siRNAs unless specified. The sequences of the siRNAs used in this study are: siBub1, CCCAUUUGCCAGCUCAAGCTT; siCdk11 #1, AGCGGCUGAAGAUGGAGAA; siCdk11 #6, GAGCGAGCAGCAGCGUGUGUU; siCdk11 #2, GAUGAAAUUGUGGCUCUAA (MU-004687-02, Dharmacon); siCdk11 #3, UGAAACACCUGCACGACAA (MU-004687-03, Dharmacon); siCdk11 #4, UAAAGCGGCUGAAGAUGGA (MU-004687-04, Dharmacon); siCdk11 #5, CAGAUGAAAUUGUGGCUCU (MU-004687-05, Dharmacon).

### Antibodies, Immunoblotting, and Immunoprecipitation

Antibodies used in this study were listed in the following: anti-centromere antibody (ACA or CREST-ImmunoVision, HCT-0100), anti-Actin (Invitrogen, MA5-11869), anti-GFP (Abcam, ab1218), anti-Myc (Roche, 11667203001), anti-Smc1 (Bethy, A300-055A), anti-Rpb1 (Abcam, 4H8, ab5408), anti-Rpb1-pSer2 (Biolegend, H5; ActiveMotif, 61083), anti-Cdk11 (Bethy, A300-310A). Anti-APC2, anti-Sgo1, and anti-Bub1 antibodies were made in-house as described previously (Liu et al., 2015). Anti-Sororin antibody was a gift from Dr. Susannah Rankin. For immunoblotting, the secondary antibodies were purchased from Li-COR: IRDye^®^ 680RD Goat anti-Mouse IgG Secondary Antibody (926-68070) and Goat anti-Rabbit IgG Secondary Antibody (926-32211).

Immunoprecipitation was performed as follows. Cells were cross-linked with buffer (50 mM Hepes, pH 8.0, 1% formaldehyde, 100 mM NaCl, 1 mM EDTA, and 0.5 mM EGTA) at room temperature for 10 min and further treated with 125 mM Glycine for another 5 min. Cells were resuspended in lysis buffer (25 mM Tris-HCl at pH 7.5, 50 mM NaCl, 5 mM MgCl_2_, 0.1% NP-40, 1 mM DTT, 0.5 μM okadaic acid, 5 mM NaF, 0.3 mM Na_3_VO_4_ and 100 units ml^-1^ Turbo-nuclease (Accelagen)). After a 1-hr incubation on ice and then a 10-min incubation at 37 °C, the lysate was cleared by centrifugation for 15 min at 4 °C at 20,817g. The supernatant was incubated with the antibody beads overnight at 4 °C. The beads were washed four times with wash buffer (25 mM Tris-HCl at pH 7.5, 50 mM NaCl, 5 mM MgCl_2_, 0.1% NP-40, 1 mM DTT, 0.5 μM okadaic acid, 5 mM NaF, and 0.3 mM Na_3_VO_4_). The proteins bound to the beads were dissolved in SDS sample buffer, separated by SDS-PAGE and blotted with the appropriate antibodies. For immunoblotting, primary and secondary antibodies were used at 1 μg ml^−1^ concentration.

### Immunofluorescence and chromosome spread

For regular immunostaining, cells were collected and then spun onto with a Shandon Cytospin centrifuge. Cells were immediately fixed with 4% ice-cold paraformaldehyde for 4 min, and then extracted with ice-cold PBS containing 0.2% Triton X-100 for 2 min. Cells were next washed with PBS containing 0.1% Triton X-100 and then incubated with primary antibodies (1:1000 dilution) overnight at 4 °C. After washed with PBS containing 0.1% Triton X-100, cells were incubated at room temperature for 1 h with the appropriate secondary antibodies conjugated to fluophores (Molecular Probes, 1:1000 dilution). After incubation, cells were washed again with PBS containing 0.1% Triton X-100, stained with 1 μg ml^−1^ DAPI and mounted with Vectashield. For chromosome spread, nocodazole-arrested mitotic cells were swelled in a pre-warmed hypotonic solution containing 75 mM KCl for 15 min at 37 °C and then spun onto slides with a Shandon Cytospin centrifuge. Cells were then treated the same as described in the regular staining.

The images were taken by a Nikon inverted confocal microscope (Eclipse Ti2, NIS-Elements software) with a ×60 objective. ImageJ and Adobe Photoshop Image processing were used to further process the obtained microscope images. Quantification was performed with ImageJ. Statistical analysis was carried out with GraphPad Prism.

### EU chasing and purification of EU-RNAs

Extraction of EU-RNAs was performed according to the protocol from Click-iT^TM^ Nascent RNA Cpature Kit (C10365, ThermoFisher). EU in a final concentration of 0.5 mM was added into for cells with a confluency of 60-80% in 10 cm petri dishes 1 hr before harvest. Then collected EU-chased cells were dissolved in TRIzol solution (Invitrogen, 15596026) and extracted total RNAs were dissolved in nuclease-free water. After being further treated with TURBO DNase (Invitrogen, AM2238) in the presence of RNase inhibitor (NEB, M3014) at 37 °C for 1 hr, total RNAs were then extracted with Phenol/Chloroform/Isoamyl alcohol (Invitrogen, 15593-031), precipitated with ice-cold ethanol solution containing glycogen (Roche, 34990920) and sodium acetate (Invitrogen, AM9740), and finally dissolved in nuclease-free water (Invitrogen, 10977-015). Purified total RNAs were then further incubated with streptavidin dynabeads (Invitrogen, 65602) pretreated with Salmon sperm DNA (Invitrogen, 15632011) in binding buffer for 45 min. With the help of DynaMag^TM^-2 Magnet (Invitrogen 12321D), dynabeads were washed with wash buffer I and II. Washed dynabeads were ready for later analyses.

### Reverse transcription and real-time PCR analysis

EU-RNA bound dynabeads were mixed with iScript Reverse Transcription Supermix (Bio-Rad, 1708841) and reverse transcription was carried out according to the manufacturer’s protocols Synthesized cDNAs were mixed with the SsoAdbanced^TM^ Universal SYBR® Green Supermix (Bio-Rad, 1725274) and then subjected to real-time PCR analysis using QuantStudio 6 Flex Real-Time PCR System (Applied Biosystems).

The primers for human cells were used in this study: GAPDH-F: 5’-TGATGACATCAAGAAGGTGGTGAAG-3’, GAPDH-R: 5’-TCCTTGGAGGCCATGTGGGCCAT-3’; Rpl30-F: 5’-CAAGGCAAAGCGAAATTGGT-3’, Rpl30-R: 5’-GCCCGTTCAGTCTCTTCGATT-3’; SAT-4-F: 5’-CATTCTCAGAAACTTCTTTGTGATGTG-3’, SAT-4-R: 5’-CTTCTGTCTAGTTTTTATGTGAATATA-3’; SAT-13/21-F: 5’-TAGACAGAAGCATTCTCAGAAACT-3’; SAT-13/21-R: 5’-TCCCGCTTCCAACGAAATCCTCCAAAC-3’. Intergenetic-1: CCAAACTTGCTTACTCCAAAGC; ATTCCAACCCAGAACCCAGA. Intergenetic-2: GGCAAATAGTCAACTTTCACTGC; CTATGGGAGGGTTGCTTTGA. Both the Intergenetic primers were described in (Khaitovich et al., 2006)

### Chromatin immunoprecipitation (ChIP)

Cells were firstly cross-linked by formaldehyde with buffer (50mM Hepes, pH 8.0, 1% formaldehyde, 100 mM NaCl, 1mM EDTA, and 0.5 mM EGTA) at room temperature for 10 min and further treated with 125 mM glycine for another 5 min. After being resuspended in immunoprecipitation buffer (10 mM Tris 8.0, 300mM NaCl, 1 mM EDTA, 1 mM EGTA, 1% Triton X-100, and 1% sodium deoxycholate), cells were sonicated using a Thermo Fisher Scientific sonicator. Cell debris was removed centrifugation and the supernatant was precleared with protein-A beads (Santa Cruz; SC-2001) at 4°C for 2 h. Precleared cell lysates were then incubated with 5 µg anti-Myc or RNAPII-pSer2 antibodies overnight and further with protein-A beads for another 2 h at 4°C. Pelleted beads were sequentially washed by low salt buffer (20 mM Tris 8.0, 150 mM NaCl, 0.1% SDS, 1% Triton X-100, and 2 mM EDTA), high-salt buffer (20 mM Tris 8.0, 500 mM NaCl, 0.1% SDS, 1% Triton X-100, and 2 mM EDTA), LiCl buffer (10 mM Tris 8.0, 0.25 M LiCl, 1% IGEPAL CA630, 1% sodium deoxycholate, and 1 mM EDTA), and TE buffer (10 mM Tris, pH 8.0, and 1 mM EDTA, pH 8.0). Afterwards, beads were treated with elution buffer (10 mM Tris 8.0, 1 mM EDTA, and 1% SDS) at 65°C for 10 min, and the elute was further incubated at 65°C overnight to reverse the cross-linking. Then the solution was sequentially treated with RNase A (Qiagen; 1007885) at 37 °C for 1 h and Proteinase K (Thermo Fisher Scientific; EO0491) at 50°C for 2 h. Finally, DNA in the solution was extracted with Phenol/Chloroform/Isoamyl alcohol (25:24:1, vol/vol; Invitrogen; 15593-031) and purified by Qiagen gel purification kit for later analyses.

### Quantification and Statistical analysis

Microscope images were imported into Image J. In the experiments of **Figures 1B, 1E, 2B, 3B, 3D, 3F, 4E, 4G, 7C, S1C, S2C, and S3C**, Five or six kinetochores were randomly selected from each cell. A mask was generated to mark centromeres on the basis of ACA fluorescence signals in the projected image. Numeric intensities of these marked signals were obtained. After background subtraction, the intensities of Bub1, Sgo1, Rpb1, Rpb1-Ser2, p-MELT, and ACA signals within the mask were obtained in number. Relative intensity was calculated from the intensity of Bub1, Sgo1, Rpb1, Rpb1-Ser2, or p-MELT signals normalized to the one of ACA signals and plotted with the GraphPad Prizm software.

For quantification of Rpb1 levels on chromosome arms in **Figure 4H**, a mask was generated to mark a chromosome arm. Numeric intensities of these marked signals were obtained. After background subtraction, the intensities of Rpb1 and DAPI fluorescence signals within the mask were obtained in number. Relative intensity was derived from the intensity of Rpb1 normalized to the one of DAPI signals and plotted with the GraphPad Prism software.

Measurement of sister-centromere distance in **Figures 1C, 2D, 3G, 4C, 7D, 7F, S2A and S3B** was carried out with Image J. A straight line was drawn between a pair of sister centromeres, revealed by ACA signals. Numeric values were automatically generated by Image J.

Quantification was usually performed based on the results from a single experiment or three independent experiments, which is specified in the legends of each experiment. Differences were assessed using ANOVA followed by pairwise comparisons using Tukey’s test. All the samples analyzed were included in quantification. Sample size was recorded in figures and their corresponding legends. No specific statistical methods were used to estimate sample size. No methods were used to determine whether the data met assumptions of the statistical approach.

**Source data files for all the related figures can be found in the fold “Source data” of the supplemental**

## Acknowledgements

This work was supported by Tulane University startup funds and the National Institute of General Medical Sciences/the National Institutes of Health grants R01GM124018 and R01GM141123, awarded to H. Liu. Research reported in this study was also supported by the National Cancer Institute of the National Institutes of Health under Award Number R01CA261258 to Z. Lin. The content is solely the responsibility of the authors and does not necessarily represent the official views of the National Institutes of Health.

## Author Contributions

Author contributions: Conceptualization, H. Liu; Methodology, Q. Zhang, YJ. Chen, Z. Teng, Z. Lin and H. Liu; Investigation, Q. Zhang, YJ. Chen, and Z. Teng; Writing-original draft, Q. Zhang and H. Liu; Writing-review and editing, Q. Zhang, YJ. Chen, Z. Teng, and H. Liu; Funding acquisition, H. Liu; Resources, H. Liu; Supervision, H. Liu.

---

**Figure S1.**
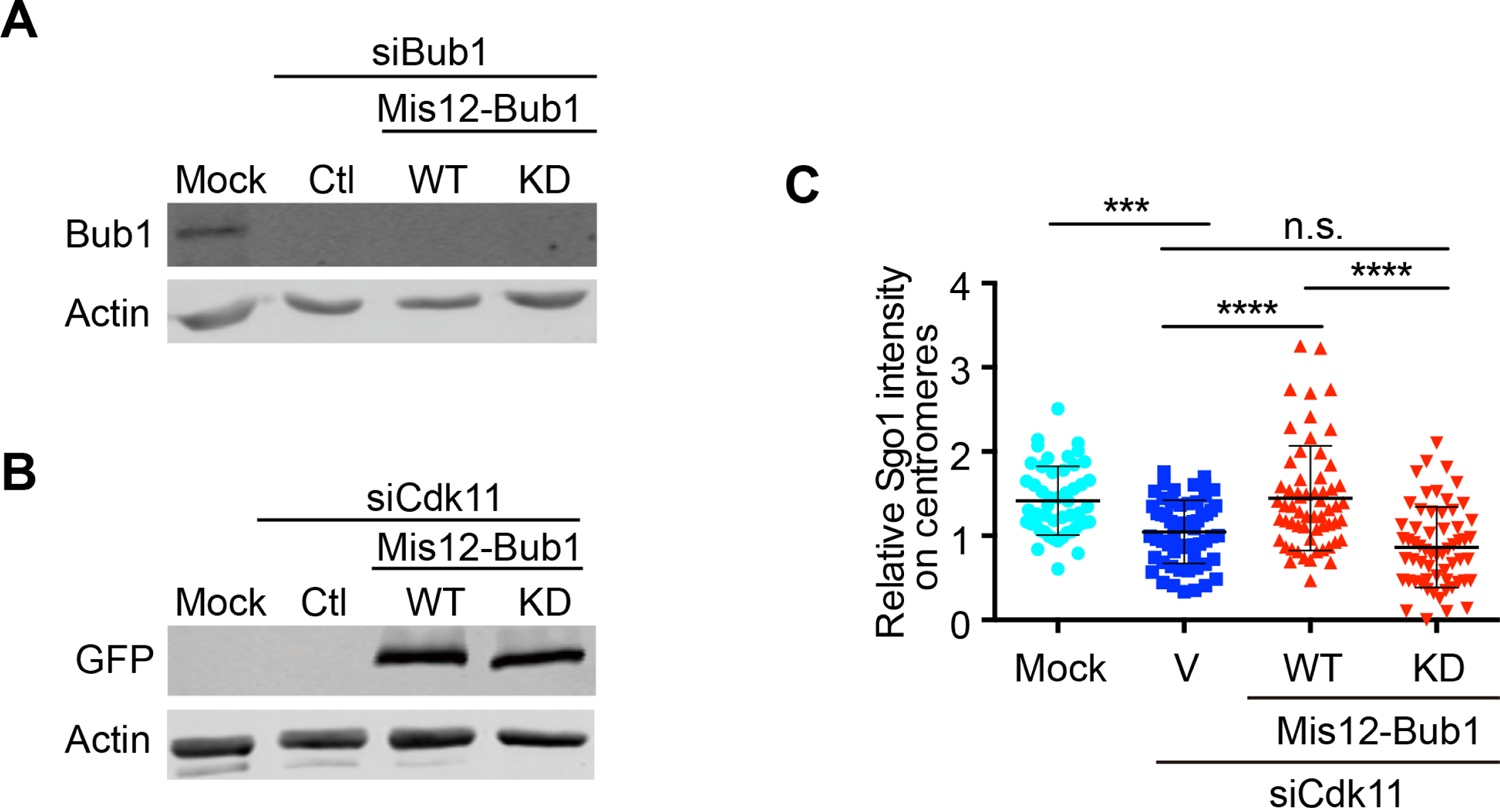
**A** and **B**. Cell lysates in (Figures 2A and 2C) were resolved with SDS-PAGE and blotted with the indicated antibodies. **C.** Quantification of relative Sgo1 levels on centromeres (Sgo1/ACA) in (Figure 2C). The quantification was performed based on a single experiment. At least 50 centromeres (5 per cell) were scored for each condition. The average and standard deviation are shown here. The experiment was repeated three times and the results were highly reproducible. n.s. denotes not significant; **, P<0.01; ****, P<0.0001.

**Figure S2.**
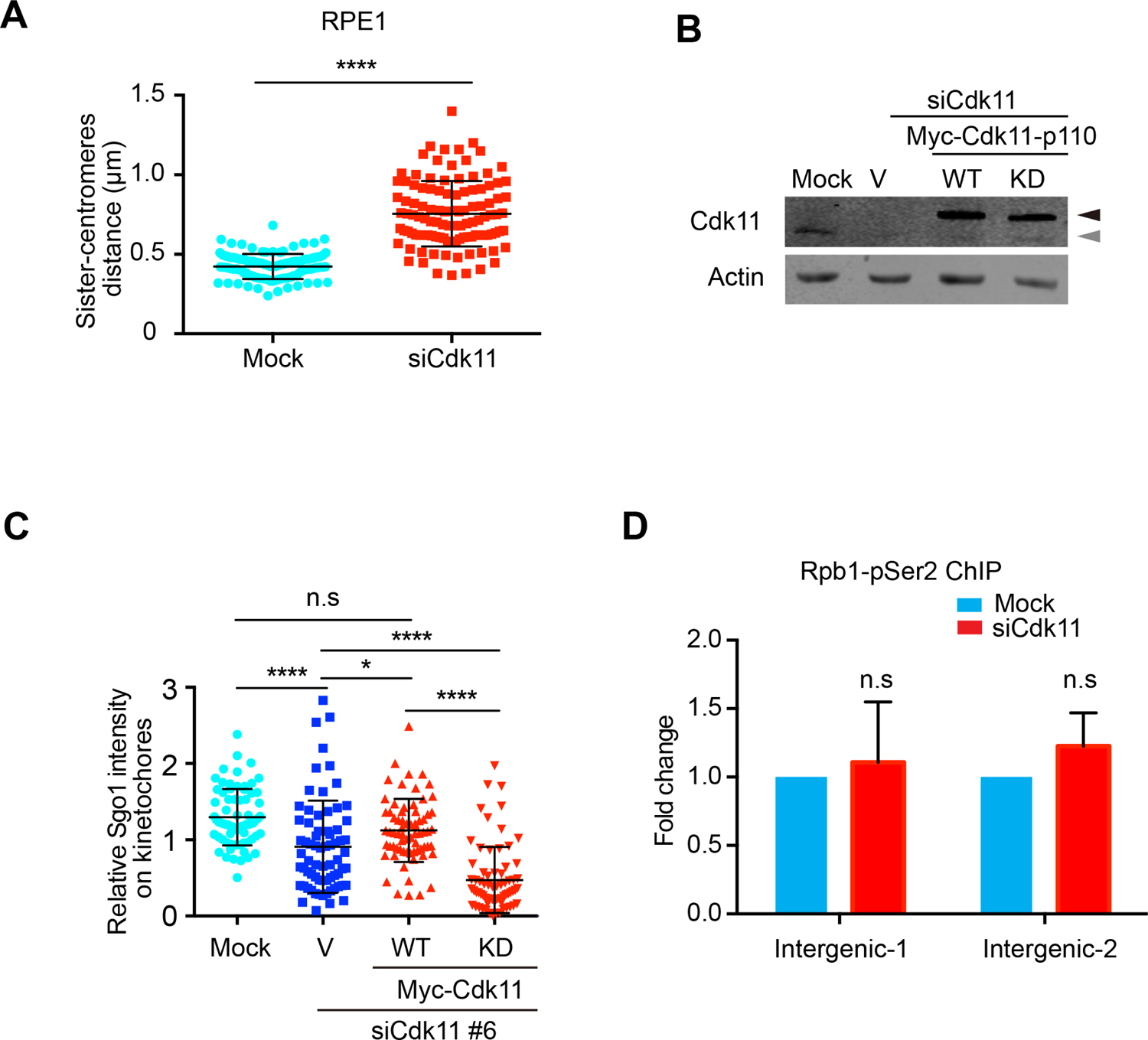
**A.** Quantification of the distance of inter-sister centromeres in (Figure 3C). The quantification was performed based on a single experiment. At least 110 sister-centromere pairs (10 per cell) were scored for each condition. The average and standard deviation are shown here. The experiment was repeated twice and the results were highly reproducible. **B.** Cell lysates in (Figure 3E) were resolved with SDS-PAGE and blotted with the indicated antibodies. Dark and grey arrowheads indicate Myc-tagged and endogenous Cdk11-p110, respectively. **C.** Quantification of relative Sgo1 levels on centromeres (Sgo1/ACA) in (Figure 3E). The quantification was performed based on a single experiment. At least 60 centromeres (5 per cell) were scored for each condition. The experiment was repeated three times and the results were highly reproducible. n.s. denotes not significant; *, P<0.05; ****, P<0.0001. **D.** Cdk11 depletion has no effects on RNAPII pSer2 association with intergenic regions. Two pairs of primers targeting Intergenic regions were applied to detect the association of RNAPII pSer2 using the samples from (Figure 3H). The average of normalized fold changes (IP/Input) and stand error calculated from three independent experiments are shown here. n.s. denotes not significant.

**Figure S3.**
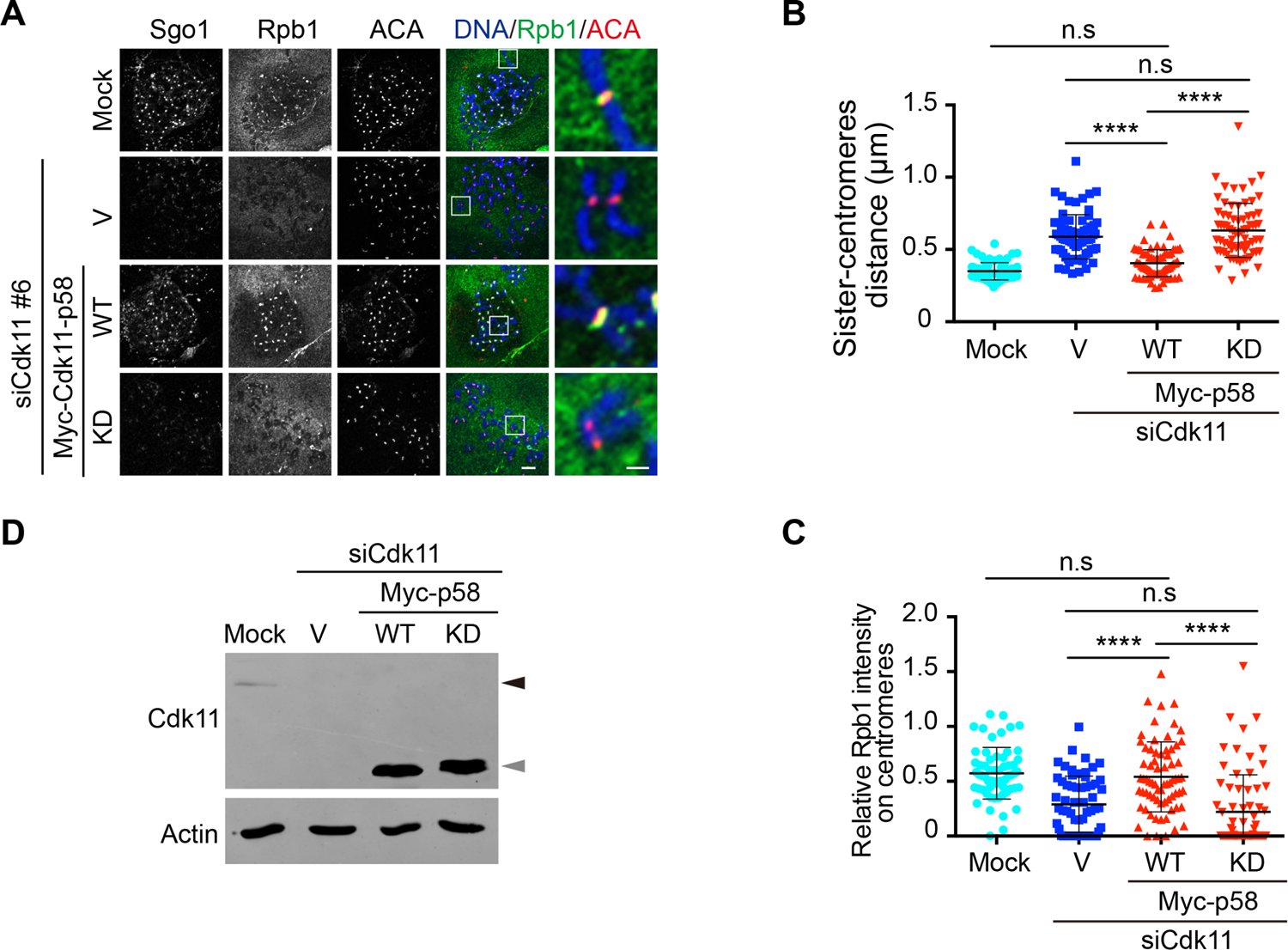
Expression of Myc-Cdk11-p58 WT, not KD, rescues the defects caused by Cdk11 depletion. **A.** HeLa Tet-On cells were transfected with vectors or plasmids containing Myc-Cdk11 WT or KD. Cells were then treated with nocodazole for 2 hr and mitotic cells were collected for chromosome spread and immunostaining with the indicated antibodies. Scale bars, 5 μm and 1 μm, respectively. **B** and **C**. Quantifications of relative Rpb1 levels (Rpb1/ACA, **B**) and the distance of inter-sister centromeres (**C**) in (**A**). These quantifications were performed based on a single experiment. At least 60 centromeres (5 per cell) were scored for each condition in (**B**) and at least 70 sister-centromere pairs (7 per cell) were scored for each condition in (**C**). The average and standard deviation are shown here. The experiment was repeated twice and the results were highly reproducible. **D.** Cell lysates in (**A**) were resolved with SDS-PAGE and blotted with the indicated antibodies. Dark and grey arrowheads indicate endogenous Cdk11-p110 and Myc-tagged G2/M Cdk11-p58, respectively.

**Figure S4.**
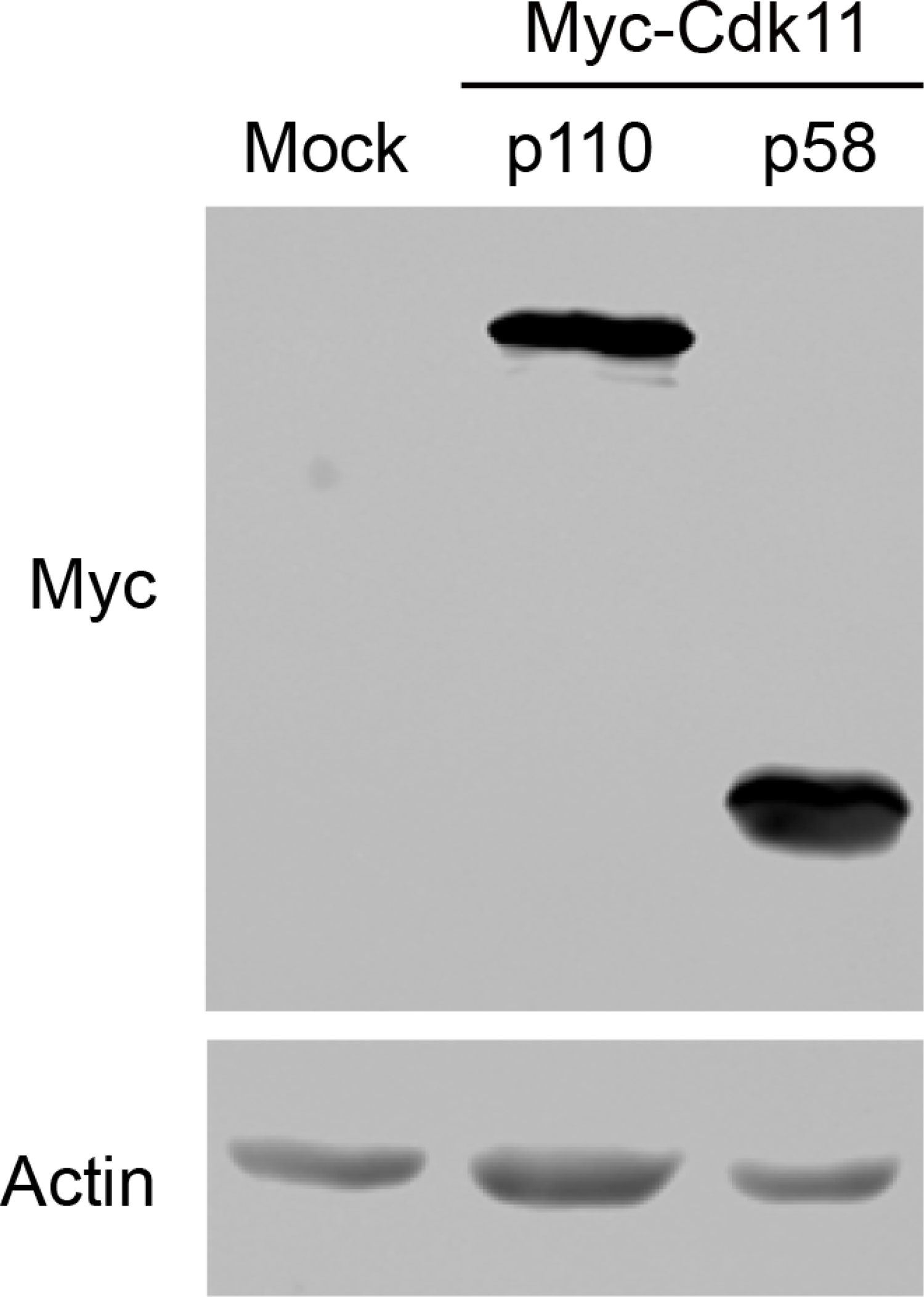
Cell lysates in (Figure 4H) were resolved with SDS-PAGE and blotted with the indicated antibodies.

**Figure S5.**
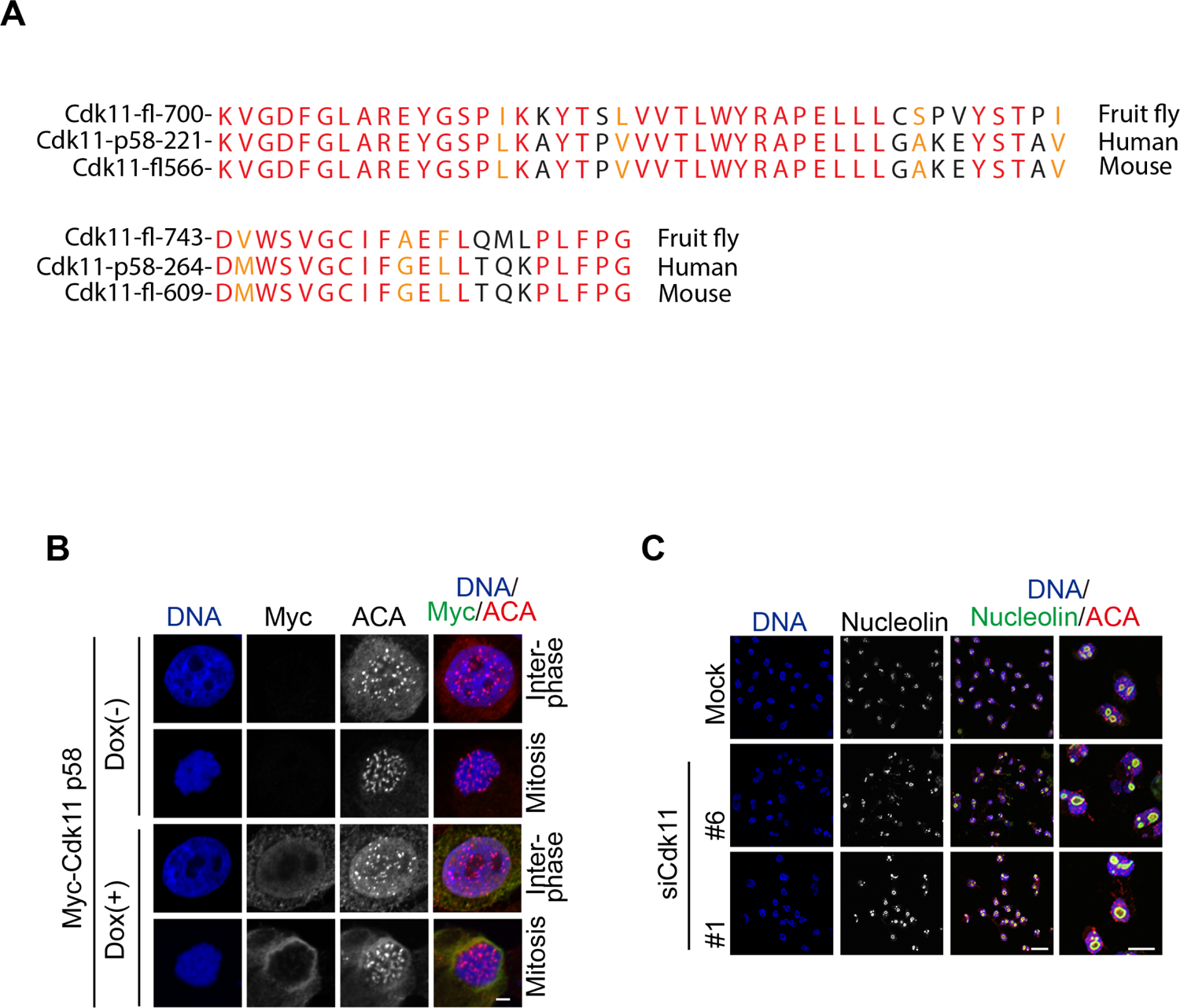
Localization of Cdk11-p58 and Cdk11 impact on nucleolar structure. **A.** The residues 221-284 in human Cdk11-p58 (corresponding to 576-640 in Cdk11-p110) are highly conserved across three species. Highly conserved amino acids are marked in red and similar amino acids in orange. fl: denotes full length. Human Cdk11-p110 kinase domain: residues 441-723. **B.** Inducible HeLa Tet-On stable cells with Myc-Cdk11-p58 were treated with or without doxycycline. Log-phase cells were then treated with nocodazole for 2 hr and mitotic cells were collected for immunostaining with the indicated antibodies. Scale bar, 5 μm. **C.** HeLa Tet-On cells were transfected with mock or Cdk11 siRNA. Log-phase cells were collected for immunostaining with the indicated antibodies. Scale bars, 20 μm and 10 μm, respectively.

**Figure S6.**
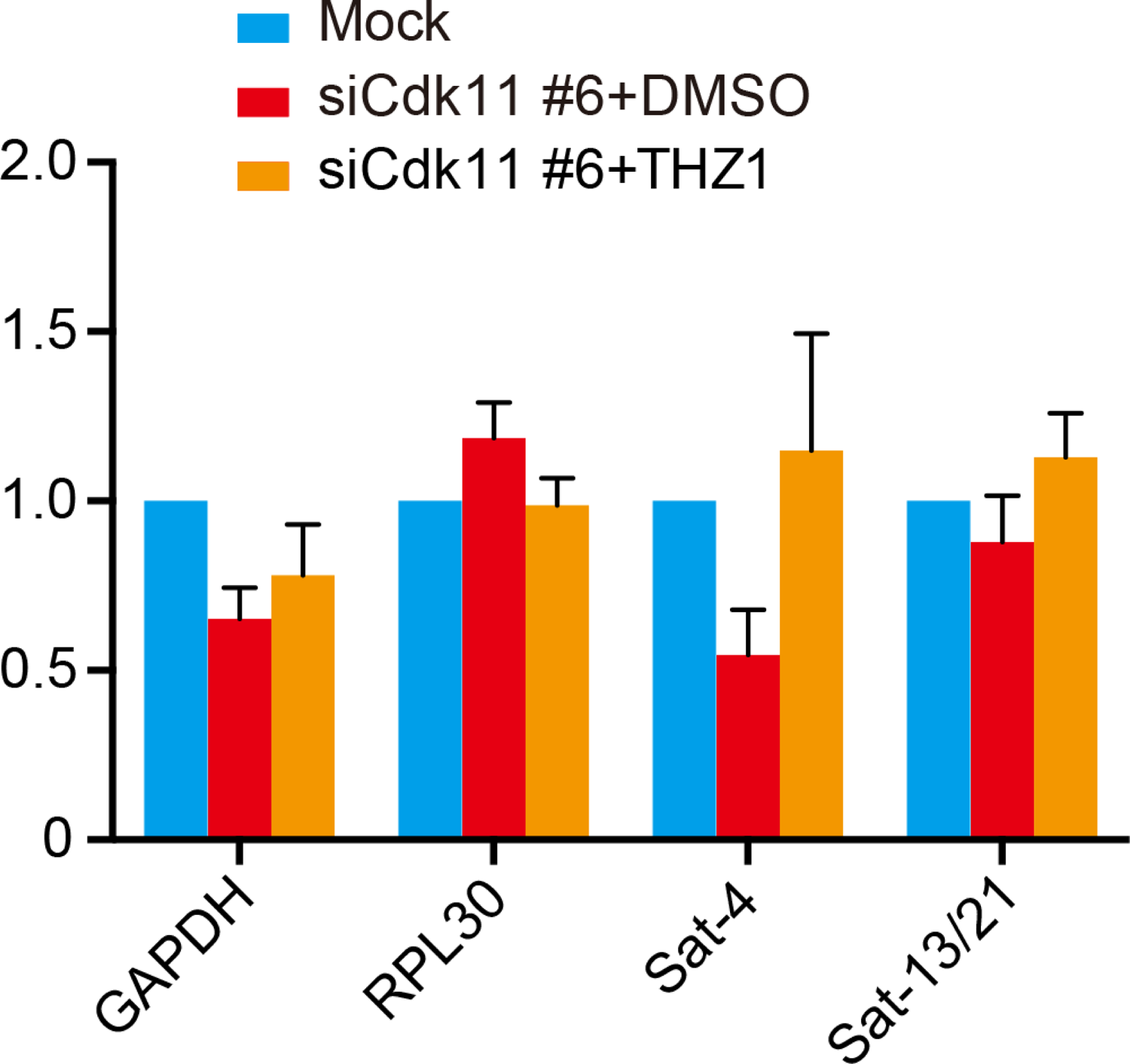
Treatment of THZ1 restores the amount of centromeric alpha-satellite RNAs in Cdk11 depletion cells Log-phase HeLa Tet-On cells transfected with mock or Cdk11 siRNAs (#6) were treated with DMSO or THZ1 for 12 hr and total RNAs were extracted for real-time PCR analysis. The average and stand error calculated from five independent experiments are shown here.

